# Vagal afferent activation induces a NREM-sleep-like state with brain-body cooling, but locus coeruleus activation

**DOI:** 10.1101/2025.03.14.643122

**Authors:** Najma Cherrad, Georgios Foustoukos, Laura MJ Fernandez, Alejandro Osorio-Forero, Yann Emmenegger, Paul Franken, Anita Lüthi

## Abstract

Parasympathetic nervous system activity, vital for restorative sleep, is influenced by vagal sensory afferents, but how these shape brain-body correlates of sleep remains open. We investigated this by combining vagal sensory neuron (VSN) stimulation with EEG/EMG, heart rate, brain-body temperature, and noradrenergic *locus coeruleus* (LC) neuron population activity measures. Opto-/chemogenetic VSN manipulations in Vglut2-Cre mice were validated via *in vitro* synaptic physiology, cFos expression, and heart rate measures. Chemogenetic VSN stimulation during the resting phase induced a non-rapid-eye-movement sleep (NREMS)-like state with enriched, but homeostatically regulated, low-frequency (0.75–4 Hz) EEG activity, yet suppressed REMS. Simultaneously, brain-body temperatures dropped while LC activity increased. Recovery was gradual towards normal NREMS-REMS-cycles but accelerated by external warming, identifying cooling as key factor. We conclude that VSN activation can promote brain-body correlates as a continuum of normal NREM sleep while engaging forebrain neuromodulation, offering insights into mechanisms of sleep disruptions linked to autonomic dysregulation.

## Introduction

The transition from wake to sleep involves a complex autonomic (de Zambotti et al., 2018) and thermoregulatory (Krauchi & Deboer, 2010) transformation that is vital for cardiovascular (Silvani et al., 2015) and gastrointestinal (Orr et al., 2020) health. In turn, sleep is negatively affected by cardiovascular and gastrointestinal diseases, which can lead to a cycle of worsening conditions (de Zambotti et al., 2018; Kourbanova et al., 2022; Miglis, 2016; Orr et al., 2020; Wei, 2019). Although the interplay between the sleeping brain and the body has long been recognized, the role of peripheral signals in sleep state regulation remains little understood.

Common experience tells us that sensations from the body influence sleep (Wei, 2019). Some of these interactions are sleep-disruptive (Bastuji et al., 2008), while others may improve sleep. For example, gastrointestinal distention can induce low-frequency brain activity that is typical for non-rapid-eye-movement sleep (NREMS) (for review, see (Orr & Chen, 2005)). Activation of carotid sinus baroreceptors can increase or decrease arousal levels depending on the intensity of stimulation (for review, see (Silvani et al., 2015)), and optogenetic activation of barosensitive brainstem neurons was reported to promote NREMS (Yao et al., 2022), while broadband decreases in EEG power were observed upon stimulation of vagal sensory neurons (VSNs) innervating the cardiac ventricular wall (Lovelace et al., 2023). Vagal sensory afferents hence supply the brain with a rich source of signals that could modify sleep. However, it is unclear whether their activity is linked to specific spectral signatures of the sleep electroencephalogram (EEG) or impacts sleep architecture and its autonomic and thermoregulatory correlates more extensively.

A large portion of sensory information from the body is relayed to the brainstem via the vagus nerve that is primarily composed (∼80 %) of chemo-, mechano-, osmo- and temperature-sensitive fibers arising from cardiovascular, gastrointestinal and respiratory systems (Berthoud & Neuhuber, 2000; Karemaker, 2022; Prescott & Liberles, 2022). The VSNs are bilaterally located in the jugular-nodose ganglia (JNG) that target the medullar *nucleus tractus solitarius* (NTS) and the *area postrema* (AP) as part of the vagal-recipient dorsal vagal complex (DVC) (Prescott & Liberles, 2022). Both the NTS and the AP project to the autonomic motor nuclei of the DVC, including the parasympathetic dorsal motor nucleus of the vagus (Neuhuber & Berthoud, 2021; Zhang et al., 2021), but also to brainstem, midbrain and hypothalamic circuits involved in sleep-wake control, including the monoaminergic *locus coeruleus* (LC) and dorsal raphe nuclei, the parabrachial nucleus, and preoptic hypothalamic areas (Holt, 2022). Activation of monoaminergic and cholinergic arousal systems by vagus nerve stimulation in rodents is particularly well documented and supported in humans through pupillometry (Berger et al., 2021). The degree of VSN activity could thus modulate sleep in diverse ways, including through its actions on both the autonomic nervous system and brain circuits of arousal regulation. However, a comprehensive characterization of the actions of specific activation of VSNs on sleep is yet to be conducted.

To investigate the impact of VSN activation on sleep, we specifically activated VSNs in sleeping mice at the onset of their preferred resting phase through chemogenetic stimulation of virally transduced JNG sensory neurons. This induced a dose-dependent expression of a prolonged (tens of minutes to hours) NREMS-like state with spectral properties distinct from, yet gradually reverting to, physiological NREMS. While this state was interrupted by numerous short arousals, REMS did not appear but recovered in the subsequent 24 h. The sleep alterations, including the delay to REMS re-appearance, correlated tightly with a brain-body temperature drop. At the same time, noradrenergic LC neuron activity became markedly elevated. The recovery to normal NREMS-REMS cycles was accelerated when we counteracted brain-body cooling by ambient warming. Together, we show that VSN activation generates a so far undescribed brain-body state that is strengthened along a continuum of spectral, autonomic and thermoregulatory patterns typical for NREMS. However, these features coincide with a noradrenergic activity that shows wake-like activity patterns. Thus, while brain-body communication is commonly thought to align physiological conditions within the organism, our data show that a sedated body can coexist with brain signatures of wakefulness.

## Results

### Vglut2-expressing VSNs target *nucleus tractus solitarius* (NTS) neurons through glutamatergic synapses

The majority of VSNs express the vesicular glutamate transporter type 2 (Vglut2), which distinguishes them from cholinergic vagal motor neurons (Chang et al., 2015). In whole-mounts of extracted left JNG (L-JNG) immunostained for VGLUT2 protein, we observed multiple fluorescently labeled cell bodies located within the ganglion and lining its borders, while sparing the sites of passage of motor fibers (**Figure 1a**). We transfected Vglut2-Cre mice with a conditional mCherry-expressing viral construct into the L-JNG (ssAAV8/2-hSyn1-dlox-mCherry(rev)-dlox-WPRE-hGHp(A)) to specifically target VSNs and to examine their sites of projection (**Figure 1b**, **Supplementary Figure 1**). After at least 3 weeks post-transfection, we observed clusters of red fibers mainly entering the ipsilateral left NTS from its lateral and ventrolateral sides. Fibers were principally located within the central portion of NTS and across its antero-posterior extent (**Figure 1c**, from Bregma (Br) -7.76 mm to Br -6.7 mm, n = 3 mice), demonstrating that transfected L-JNG neurons projected to a major portion of the DVC.

**Figure 1.**
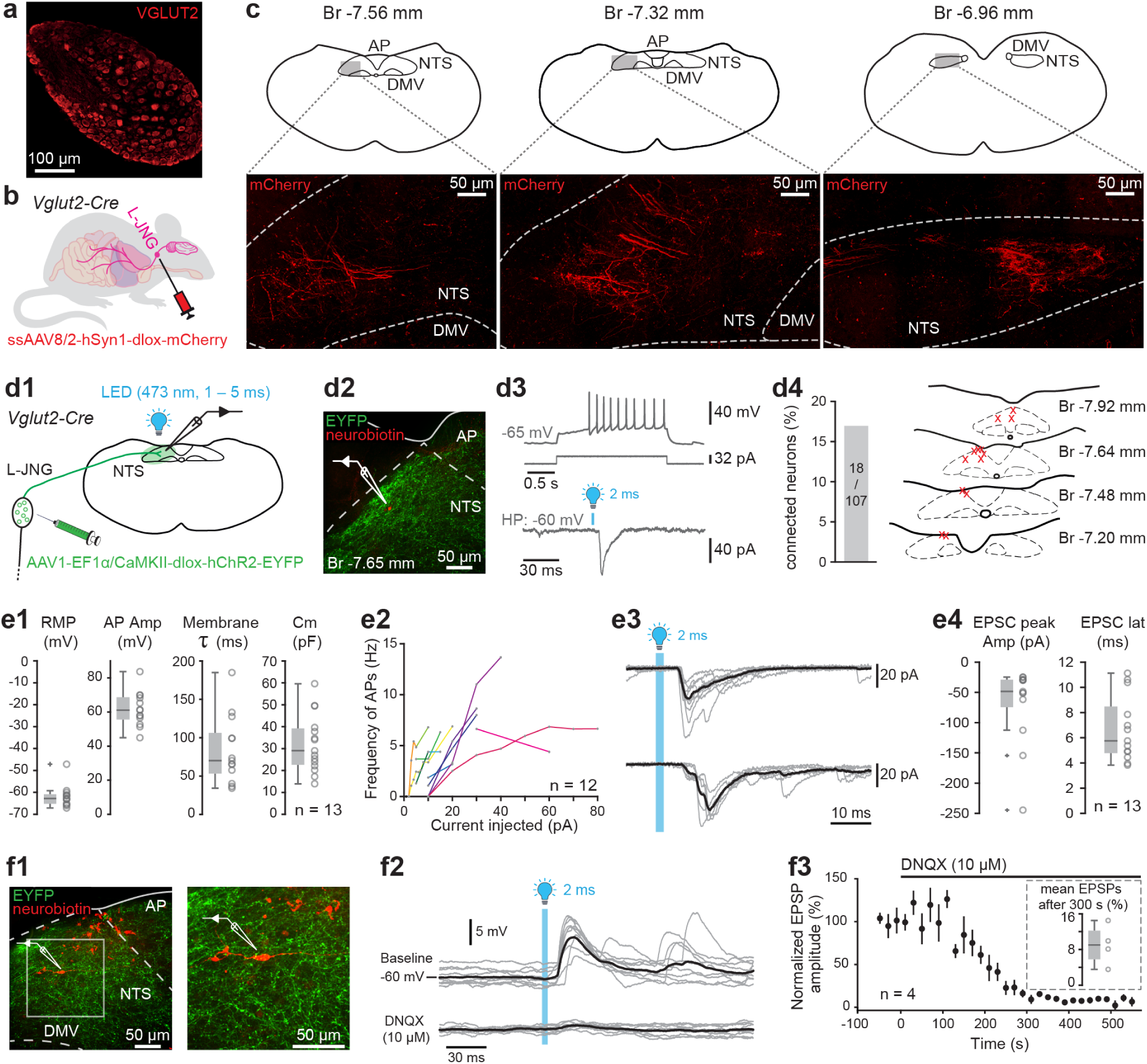
Anatomical and *in vitro* functional identification of vagal sensory neurons (VSNs) a. Fluorescence microscopy image of a left jugular nodose ganglion (L-JNG) immunostained for VGLUT2 protein. Labelled cell bodies are found throughout the ganglion, except in the area through which the axons from the motor nerve pass (top left rectangular black area within the L-JNG). b. Schematic of the viral transfection to target VSNs. The L-JNG of Vglut2-Cre mice was virally transfected with a conditional mCherry-reporter construct (ssAAV8/2-hSyn1-dlox-mCherry(rev)-dlox-WPRE-hGHp(A)). See **Supplementary Figure 1** for illustration of the surgical procedure. c. Fluorescence microscopy images of mCherry-expressing fibers in brainstem vagal-recipient areas at three different anteroposterior locations (schematic sections from mouse atlas shown on top). NTS, *nucleus tractus solitarius*, AP, *area postrema*, DMV, dorsal motor nucleus of the vagus. d. (d1) Schematic brain slice prepared for *in vitro* patch-clamp recordings from Vglut2-Cre animals virally transfected for conditional ChR2 expression, with injections of AAV1-EF1α-DIO-hChR2(H134R)_EYFP-WPRE-hGH or AAV1-CaMKIIα-hChR2(H134R)_EYFP-WPRE-hGH). Patched neurons within EYFP-labeled (green) regions were exposed to blue light pulses (see blue text for details) through the objective. (d2) Micrograph of a *post hoc* recovered NTS cell soma filled with neurobiotin (red) and surrounded by ChR2_EYFP-expressing VSN axons (green). (d3) Representative action potential firing pattern of the NTS cell shown in d2 in response to a depolarizing somatic current pulse (top two traces) and example optogenetically evoked excitatory postsynaptic current (EPSC) in the same cell voltage-clamped at a holding potential (HP) of -60 mV. (d4) Percentage of synaptically connected NTS neurons shown as bargraph, together with the localization of the 13 neurons that could be morphologically identified at different anteroposterior Bregma (Br) levels. All cell bodies were located within the NTS (dotted lines). e. (e1) Box-and-whisker plot of the intrinsic electrophysiological properties of 13 NTS neurons responding to VSN axon optogenetic stimulation. RMP, resting membrane potential (RMP); AP amp, action potential amplitude; Membrane τ, membrane time constant; C_m_, membrane capacitance. (e2) Current-frequency response curves from 12 of the 13 recorded cells; one cell had only a single value and was not included in the graph. Frequency was calculated as the mean action potential rate per somatic current injection. (e3) Example opto-evoked EPSCs from 2 different NTS neurons. Grey, individual responses; black, mean response. The top example shows a case of fixed response latency, the bottom one a case with variable response latency. (e4) Box-and-whisker plot of optogenetically evoked EPSC characteristics. Left, EPSC negative peak amplitudes, calculated for the first peak in single- and multicomponent responses. Right, EPSC latencies, calculated from the onset of the light pulse and plotted for responses with < 0.5 ms jitter. f. (f1) Micrographs of a recorded NTS cell shown at two magnifications (red) surrounded by EYFP-expressing VSN axons. (f2) Example EPSPs from the same cell, with mean response superimposed in black. Top: baseline EPSPs; Bottom: EPSPs after bath-application of the AMPA receptor antagonist DNQX (10 μM). (f3) Time course of DNQX actions on normalized EPSPs for n = 4 recordings. Inset shows the remaining response in the presence of DNQX, calculated as the mean EPSP after 300 s, expressed in percent of the baseline EPSP.

We tested the functional connectivity of L-JNG afferent fibers through studying vagally evoked synaptic responses in NTS neurons. We prepared acute brainstem slices from Vglut2-Cre mice previously transfected for conditional expression of the excitatory opsin channelrhodopsin 2 (hChR2) in L-JNG (AAV1-EF1α-DIO-hChR2(H134R)-EYFP-WPRE-hGH or AAV1-CaMKIIα-hChR2(H134R)-eYFP-WPRE-hGH) and patched cell bodies within the NTS using neurobiotin-containing patch pipettes (**Figure 1d1**). An example recorded NTS neuron recovered *post hoc* together with microscopic fluorescent visualization of VSN axons surrounding the cell body is shown in **Figure 1d2**. Somatic current injections were used to generate action potentials, and whole-field blue light pulses (470 nm, 1 – 5 ms duration, 3.8 mW) applied to evoke synaptic responses (**Figure 1d3**). We found postsynaptic currents in ∼17% (18/107) of recorded neurons and could recover the morphology of 13 connected neurons within the caudal and intermediate NTS (Br -7.9 mm to Br -7.1 mm), of which 7 were clustered in the dorsal region of the NTS adjacent to the AP (**Figure 1d4**). These 13 neurons had a mean resting membrane potential of -61.8 ± 4.8 mV (n = 13) and responded with action potentials upon depolarizing current injection, with membrane time constants and capacitance values in line with previous reports (Kline et al., 2002) (**Figure 1e1**). Tonic action potential discharges reached rates up to ∼14 Hz (**Figure 1e2**). Evoked postsynaptic currents (EPSCs), measured in voltage-clamp at a holding potential of -60 to -70 mV, appeared as multicomponent inward currents with variable latencies, suggesting combined recruitment of mono- and polysynaptic afferents (Doyle & Andresen, 2001). Indeed, in 13/18 cells, we found a fixed latency with a jitter < 0.5 ms (range 0.09 – 0.49 ms) for all evoked responses, suggesting monosynaptic connectivity. However, in 10 of these 13 recordings, responses with a larger delay and a jitter > 1 ms were also evoked, indicating that polysynaptic circuits were recruited (**Figure 1e3**). Putative monosynaptic responses showed a mean peak amplitude of -68.5 ± 62.8 pA and mean latencies of 6.7 ± 2.5 ms (range 3.9-11.1 ms, with values > 10 ms for 3/13 recordings) (**Figure 1e4**). These values are larger than what is typical for monosynaptic connections, as also observed in previous studies on the vagus-NTS synapse (Bailey et al., 2008; Miles, 1986). In 4 recordings with a putative monosynaptic connection, bath application of DNQX (10 µM), an antagonist of glutamatergic α-amino-3-hydroxy-5-methyl-4-isoxazolepropionic acid (AMPA) receptors, suppressed evoked EPSPs to ∼9 % of the original response (**Figure 1f1**-**f3**, n = 4). Therefore, virally targeted L-JNG sensory neurons activate NTS neurons through primarily ionotropic glutamatergic synapses involving both putative mono- and, in most cases, also polysynaptic pathways.

### Chemogenetic stimulation of L-JNG VSNs activates NTS neurons *in vivo* and reduces heart rate

To test whether stimulation of transfected VSNs was functionally effective *in vivo*, we focused on the L-JNG that provides afferent information from several organs including the heart (Han et al., 2018; Min et al., 2019; Williams et al., 2016). Vglut2-Cre animals transfected with a conditional excitatory Designer-receptor-exclusively-activated-by-designer-drugs (Dreadd)- and mCherry-expressing viral vector (ssAAV8/2-hSyn1-dlox-hM3D(Gq)_mCherry-dlox-WPRE-hGH(pA) were injected > 3 weeks later with NaCl or with the chemogenetic agonist clozapine *N*-oxide (CNO) at a low dose (1.5 mg/kg i.p.) that on its own does not affect spontaneous sleep-wake behavior (Fernandez et al., 2018; Traut et al., 2023) (**Figure 2a**). We sacrificed animals 60 – 90 min later for immunohistochemical quantification of cFos protein expression levels, a marker for neuronal activity (**Figure 2b**, **c1**, **Supplementary Figure 2**). CNO but not NaCl injection led to a clear presence of cFos-expressing cell bodies across the anteroposterior extent of both the left and the right NTS (**Figure 2c2**, n = 7 for NaCl and n = 8 for CNO 1.5 mg/kg), which were surrounded by mCherry-expressing VSN axons. Strong cFos expression was also found in the AP, which regulates food intake (Borgmann et al., 2021) and can promote nausea (Engström Ruud et al., 2024; Zhang et al., 2021).

**Figure 2.**
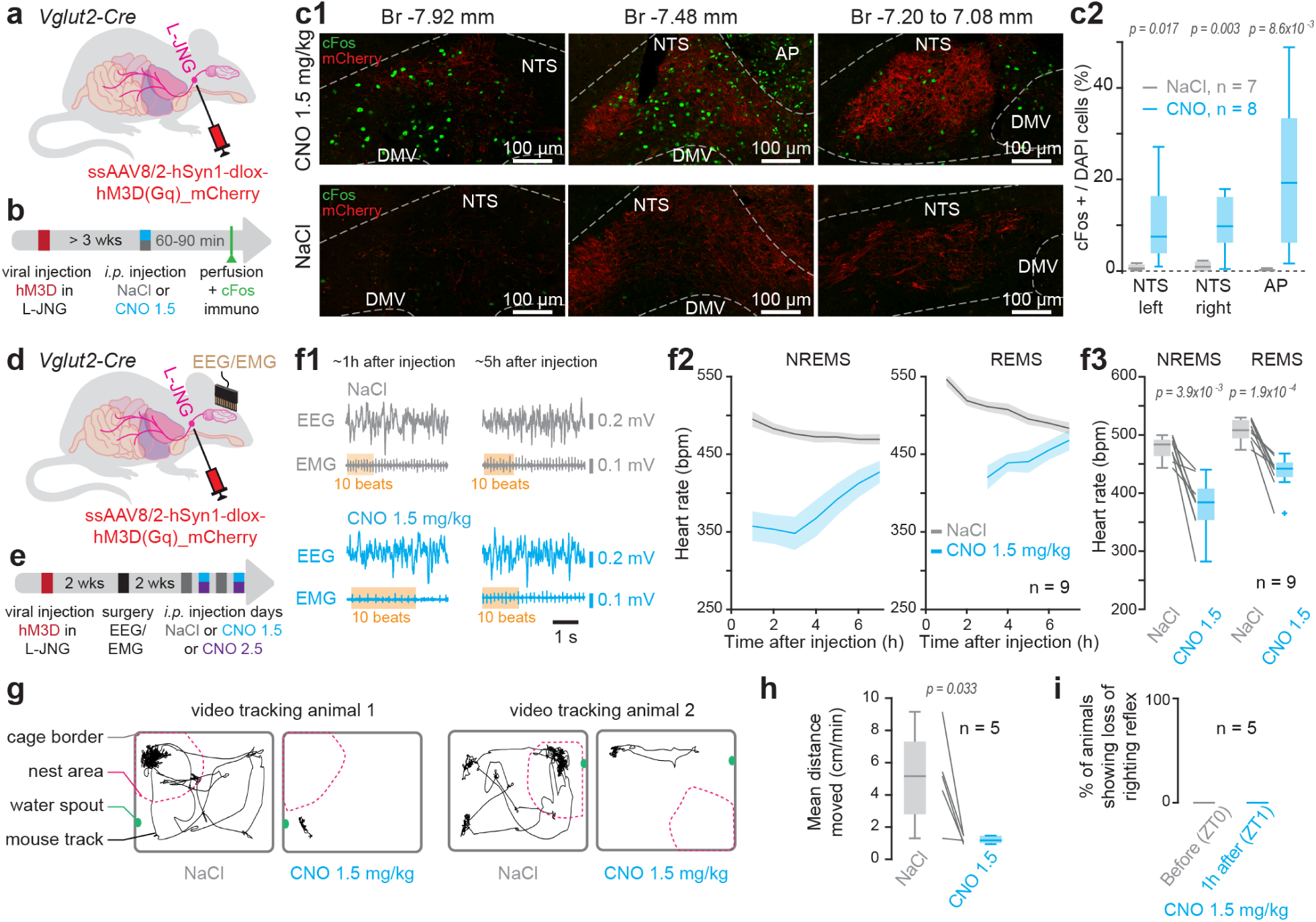
Validation of the *in vivo* efficacy of chemogenetic activation of VSNs. a. Schematic of the viral transfection surgery for chemogenetic stimulation. See **Supplementary Figure 1** for illustration of the surgical procedure. b. Experimental timeline to assess cFos expression in NTS. hM3D(Gq), excitatory chemogenetic Dreadd receptor; CNO 1.5, CNO injected at 1.5 mg/kg. c. Fluorescent micrographs for cFos expression in NTS and AP for CNO (top) and NaCl (bottom) injections. (c1), Representative examples for 3 anteroposterior Bregma (Br) levels in one mouse; (c2) quantification across animals for left and right NTS for Br -7.32 to -7.64 mm (see also **Supplementary Figure 2**). d. Schematic of the viral transfection surgery for chemogenetic stimulation in combination with EEG/EMG recordings. e. Experimental timeline. f. Heart rate in beats per min (bpm), after i.p. injection of NaCl or CNO in animals expressing hM3D(Gq). (f1) Example raw EEG/EMG signals, with the heart beats visible in the EMG traces, shown for 2 time points after i.p. injections. Colored rectangles highlight 10 heart beats. (f2) Time course of heart rates in NREMS and REMS for 5 h after injection. (f3) Quantification across animals for means across the 5 h. Wilcoxon or paired t-tests. g. Example homecage mouse trajectories during 150 min after NaCl or CNO injections. h. Quantification of mean distances moved. Paired *t*-test. i. Absence of loss of righting reflex (LORR) before and 60 min after injection of CNO 1.5 mg/kg.

We next examined heart rate to assess whether VSN activation slowed down heart rate as previously reported (Chang et al., 2015). Excitatory Dreadd-expressing animals were implanted with EEG and neck EMG electrodes for sleep-wake and heart rate monitoring (**Figure 2d**, **e**) (Fernandez et al., 2017). Upon CNO (1.5 mg/kg) injection, heart rates (quantified via the heart beats visible on the EMG trace of n = 9 /11 animals), showed a strong but reversible decrease over several hours (**Figure 2f1-f3**).

Chemogenetic VSN stimulation also affected the animals’ spontaneous locomotor behavior. Within ∼20 min after the CNO (1.5 mg/kg) injection, animals moved much less and often remained motionless outside the nest (**Figure 2g**, **h**, n = 5), suggesting profound sedation. However, these animals scored negative for the loss-of-righting reflex, showing that they remained conscious and reactive to external stimuli (**Figure 2i**, n = 5) (Gelegen et al., 2018).

### Chemogenetic stimulation of VSNs strongly increases neuronal activity in *locus coeruleus*

Vagus nerve stimulation has been reported to increase activity of monoaminergic and cholinergic arousal systems, including the noradrenergic LC (Berger et al., 2021). In line with this, we found elevated cFos expression levels in both left and right LC at 60 – 90 min after CNO (1.5 mg/kg) over NaCl injections (**Figure 3a1**, **a2**, n = 6 for for both NaCl and CNO). To explore how VSN activation modulated neuronal activity in the LC in real time, we carried out fiber photometry of bulk LC population activity while chemogenetically stimulating VSNs in combination with EEG/EMG recordings. For this purpose, we co-injected dopamine-β-hydroxylase(DBH)-(i)Cre animals with ssAAV8/2-hSyn1-hM3D(Gq)_mCherry-WPRE-hGHp(A) in the L-JNG for VSN stimulation, and with AAV5-hSyn1-dlox-jGCaMP8s-dlox-WPRE-SV40p(A) in the right LC, where we previously characterized activity patterns typical for NREMS (**Figure 3b**) (Osorio-Forero et al., 2024). Animals were then surgically implanted with EEG/EMG electrodes and, once ready for recording, injected with NaCl or CNO (1.5 mg/kg i.p.) within 30 min after light onset (Zeitgeber Time ZT = 0), the start of their predominant resting phase. We recorded EEG/EMG signals during the light phase and then gave 1 – 2 recovery days. **Figure 3c1** shows an example recording, with the hypnogram revealing the alternation between wakefulness, NREMS and REMS that is typical for mice during the light phase, together with the corresponding variations in LC activity. Control NaCl injections left these patterns unaltered, except for a brief increase shortly after the injection due to animal handling. Spontaneous LC activity patterns were instead profoundly transformed once CNO was injected, reaching steadily elevated levels that were substantially greater than any of the ones found during the spontaneous alternation between normal sleep and wakefulness (**Figure 3c2**). Furthermore, the infraslow fluctuations typical for NREMS were absent (**Supplementary Figure 3**). During this period, EEG/EMG scoring indicated that NREMS persisted and microarousals continued to be generated, while REMS was suppressed. Over a time course of several hours, which is typical for the time course of action of CNO (see e.g. (Fernandez et al., 2018)), LC activity levels returned to normal, activity fluctuations resumed and REMS re-appeared. All effects were highly consistent across animals when injected with CNO compared to NaCl (**Figure 3d1-d4**). The VSN stimulation thus dramatically elevated LC activity, producing a pattern that has so far been thought to be incompatible with states of sleep, as scored by EEG/EMG. To understand this, we undertook a detailed analysis of the properties of the sleep state generated in CNO conditions.

**Figure 3.**
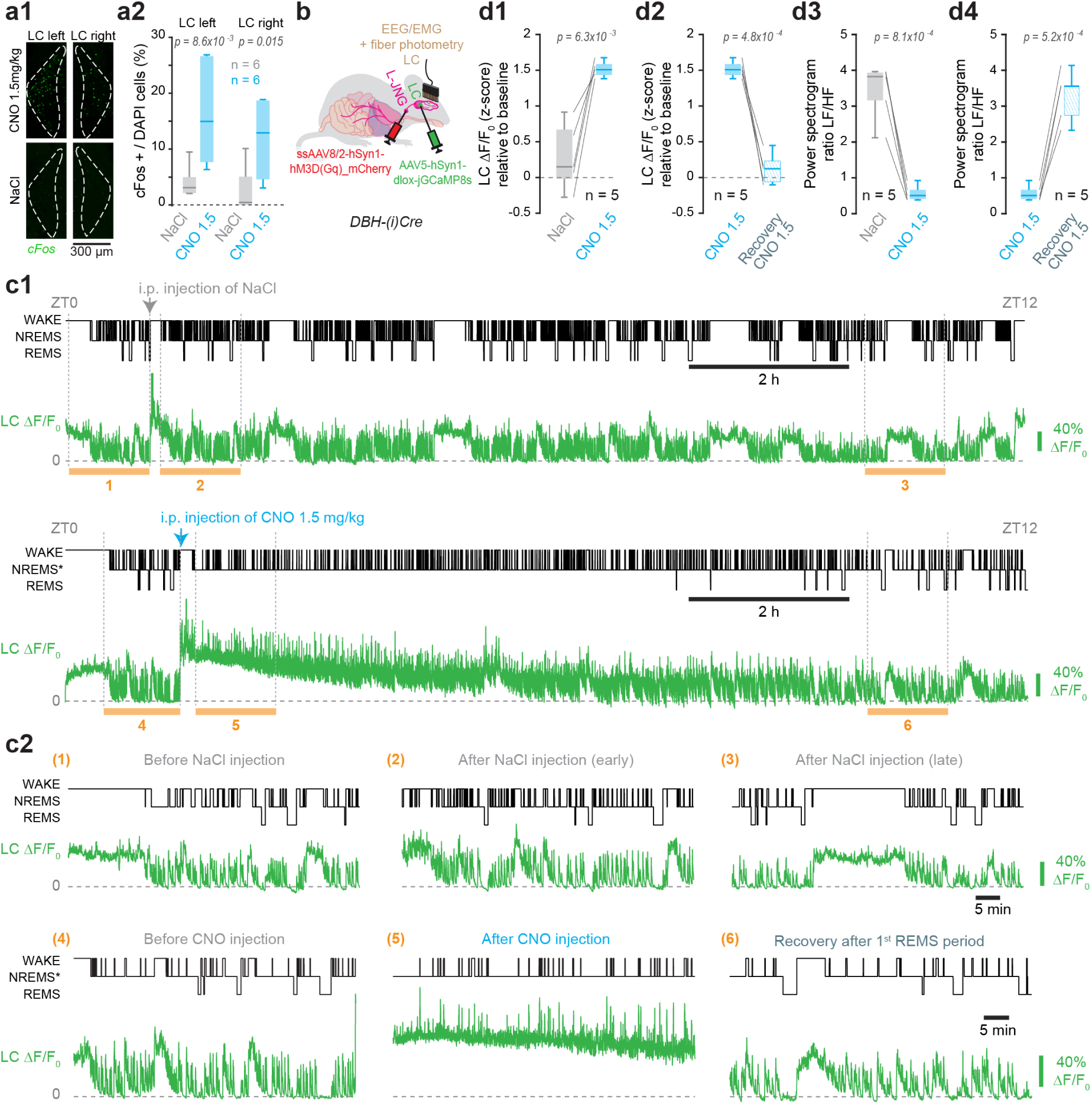
Activation of the LC by chemogenetic stimulation of vagal sensory afferents. a. (a1) Micrographs of left and right LC stained for cFos expression after CNO (top) or NaCl (bottom) injections in animals with L-JNG expressing excitatory Dreadd receptors. (a2) Box- and-whisker plot quantification for left and right LCs. Unpaired t-test (for left LC) or Wilcoxon’s rank sum (for right LC) tests with Bonferroni-corrected p = 0.025. b. Viral injection strategy for combined VSN stimulation and LC fiber photometric monitoring. DBH-(i)Cre instead of Vglut2-Cre animals were used, with a conditional construct for GcaMP8s expression injected into LC as indicated. c. Fiber photometric monitoring of bulk LC activity during chemogenetic activation of VSNs. Top, Example recording for NaCl injection showing (from top): hypnogram, ΔF/F_0_ for jGCaMP8s-derived fiber photometric signal of LC activity (green). Bottom, recording from the same animal when injected with CNO. (c2) Insets chosen from recordings in (c1), indicated by numbers. See **Supplementary Figure 3** for a spectral analysis of the LC signal. d. (d1, d2) Quantification of z-scored mean ΔF/F_0_ in LC, normalized to baseline values before injection. Means were taken for the first hour after NREMS* onset (d1) or for the first hour after the 1^st^ REMS epoch (d2, recovery). (d3, d4) As d1, d2 for power spectral activity of LC signal. The ratio of low-over high-frequency power (LF/HF) was obtained by dividing power in the lower (0.01 to 0.03 Hz) and the higher spectral components (0.1 to 0.25 Hz) to quantify the changes in infraslow activity pattern. Paired t-tests with Bonferroni-corrected p = 0.0125.

### Chemogenetic stimulation of VSNs dose-dependently induces a NREMS-like state with distinct spectral composition and overall decreased power

To assess the architecture and spectral properties of the sleep state induced by chemogenetic stimulation of VSNs, animals were injected with NaCl or CNO at two doses (1.5 or 2.5 mg/kg i.p.) within 30 min after light onset (Zeitgeber Time ZT = 0) in separate sessions interspersed by 1 – 2 recovery days. **Figure 4a1** shows an example recording from a NaCl-injected mouse, exemplifying the typical spectral dynamics for wakefulness, NREMS and REMS. Entries from wake into NREMS were accompanied by increases in EEG power in the low-frequency delta (δ, 1.5 – 4 Hz) band and the sleep spindle-containing sigma (σ, 10 – 15 Hz) band, while the ratio of the theta (θ, 6 – 8 Hz) over δ power (θ/δ ratio) marked REMS (Fernandez et al., 2018; Osorio-Forero et al., 2024). Based on these scoring rules, CNO at 1.5 mg/kg produced a spectral EEG pattern that was clearly distinct from the NaCl condition and the normal states of NREMS and REMS (**Figure 4a2**). To quantify this, we characterized the spectral composition of the EEG signals by calculating relative power spectra for the time until the first consolidated REMS bout (defined as ≥ 12 s of REMS). To compare CNO-induced spectra to their NaCl control, we used equal times spent in NREMS (n = 11). This revealed that the relative EEG δ power remained higher while σ power was decreased and did not fluctuate in a manner typical for normal NREMS (Lecci et al., 2017). Furthermore, no increases in the θ/δ ratio were detected and REMS was absent. These CNO-induced spectral alterations recovered to levels characteristic for NREMS at around 74 min after injection, a moment when REMS reappeared (**Figure 4a2**). When CNO was injected at the higher dose of 2.5 mg/kg, similar but stronger effects on sleep and the EEG were observed, and the time for recovery until the first REMS episode was longer (174 min after injection; **Figure 4a3**). Quantification across animals showed that CNO produced a brain state enriched in relative power for the low-frequency bands, notably for the slow oscillation (SO, 0.75 – 1.5 Hz) and for the δ band, while power in the σ band was reduced (**Figure 4b**, n = 11).

**Figure 4.**
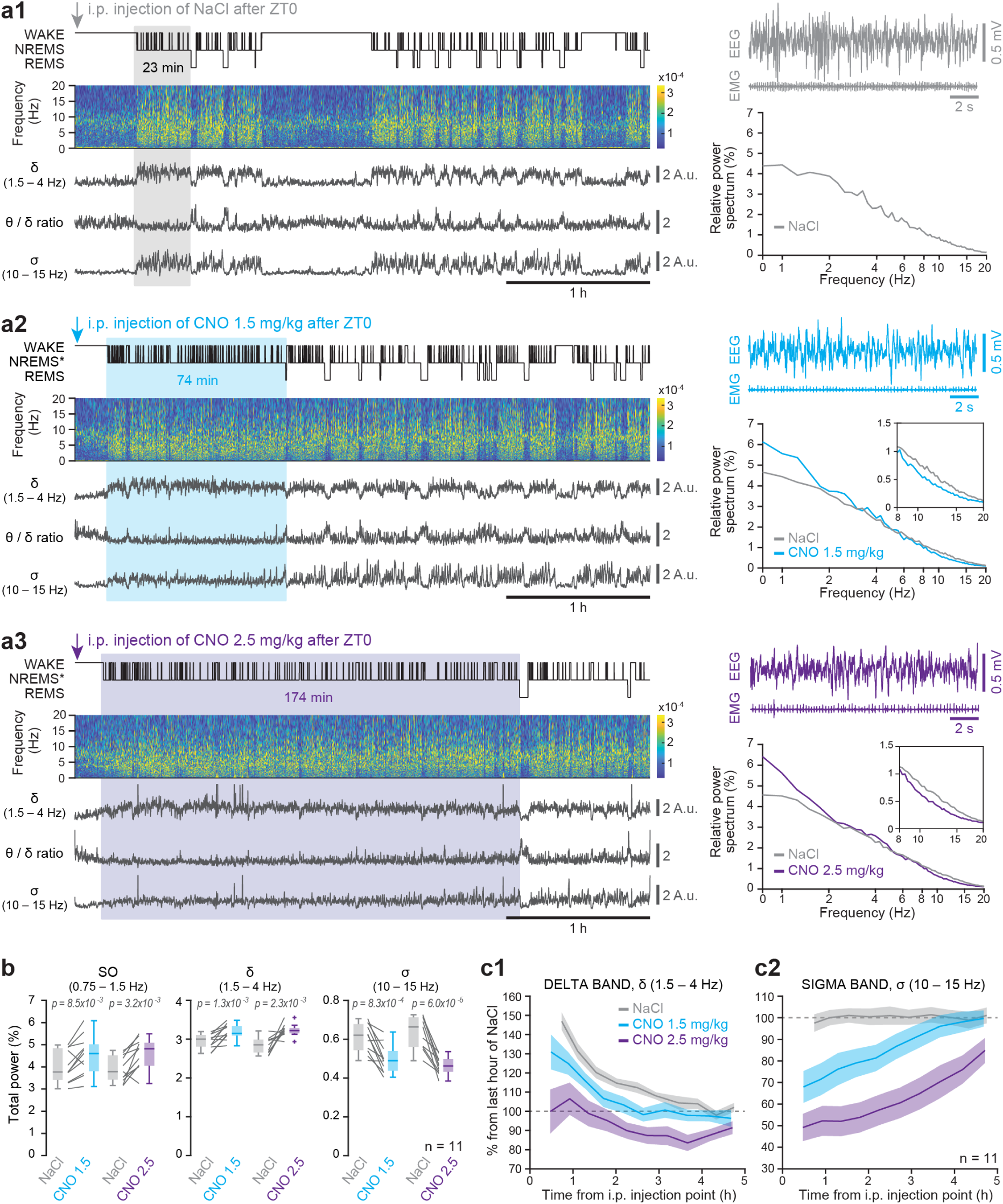
Chemogenetic activation of VSN axons induces a NREMS-like state. a. Example recordings from an excitatory Dreadd-expressing mouse, injected with NaCl (a1) or CNO at two different doses (1.5 and 2.5 mg/kg) in repeated recording sessions (a2, a3). The first 4 h of recording are shown starting at ZT0, with from top: the hypnogram, the time-frequency plot between 0 – 20 Hz, and the power dynamics of two spectral bands characteristics for NREMS, the δ band (1.5 – 4 Hz) and the σ band (10 – 15 Hz). The θ/δ ratio characterizes REMS. Colored rectangles indicate the data used to quantify the normalized relative power spectra shown on the right, together with raw EEG/EMG traces. Relative normalized power spectra, calculated by dividing by total power levels between 0.75-35 Hz for each condition, were calculated for the time until the first consolidated REMS period. Light blue, low dose of CNO (1.5 mg/kg), dark blue, high dose of CNO (2.5 mg/kg). The corresponding spectra for NaCl were calculated for the same times spent in NREMS. In a2, a3, insets show enlarged portions of the spectra to highlight the decreases in relative power levels for the σ band (10 – 15 Hz). b. Quantification of relative power in three major spectral bands typical for NREMS, the slow oscillation (SO) (0.75 – 1.5 Hz), the δ (1.5 – 4 Hz), and the σ (10 – 15 Hz) bands. Boxplots are presented for NaCl (grey), CNO 1.5 mg/kg (light blue), CNO 2.5 mg/kg (dark blue) for the three bands, with connecting grey lines showing data from individual animals (n = 11). For the two CNO conditions, corresponding NaCl values were calculated separately to match total times spent. Paired two-tailed t-tests, with significant p values indicated. c. Time course of δ (c1) and σ (c2) absolute power levels, plotted as percentage of values for the NaCl condition taken from the last two time bins (between hours 4 – 5 of the recording) to illustrate the gradual recovery of CNO to NaCl levels. See **Supplementary Figure 4** for statistical analysis.

In addition to this relative enrichment in low-over high-frequency power, EEG traces recorded in the CNO condition showed a ‘flattening’, i.e. a diminished overall amplitude (see example traces in **Figure 4a1-a3**). To verify this, we took the absolute power levels in the δ- and the σ-bands for both CNO conditions and plotted these as percentage of the NaCl values measured at the end of a 5-h interval. CNO lowered the total EEG power in both bands in a dose-dependent manner (**Figure 4c**, n = 11), but values recovered after 5 h (**Supplementary Figure 4a, b**, n = 11). Noteworthily, the spectral power changes induced by CNO recovered progressively towards the ones of NREMS. To account for this progressive transition between the CNO-induced NREMS-like state and normal NREMS, with no obvious abrupt changes to distinguish between these, we used the term ‘NREMS-like’ to refer to the CNO-induced sleep state (referred to as NREMS* in the figures).

The latency to onset of NREMS* was shorter than that for NREMS (**Figure 5a**, n = 11). As control, animals expressing a non-Dreadd-related viral construct (ssAAV8/2-hSyn1-dlox-mCherry(rev)-dlox-WPRE-hGHp(A)) showed unaltered latencies for both CNO doses (**Supplementary Figure 5**). The time spent in NREMS* was greater for the high compared to the low CNO dose and occurred at the expense of times spent in wakefulness and in REMS (**Figure 5b**, n = 11). Compared to NREMS that followed NaCl injections, NREMS* showed a preserved density of 4 – 12-sec-long microarousals from NREMS, which are characterized by EEG and EMG activation, although there was a tendency to increase with the higher CNO dose (**Figure 5c1**, n = 11). Mean bout lengths of NREMS* were also unchanged (**Figure 5c2**, n = 11).

**Figure 5.**
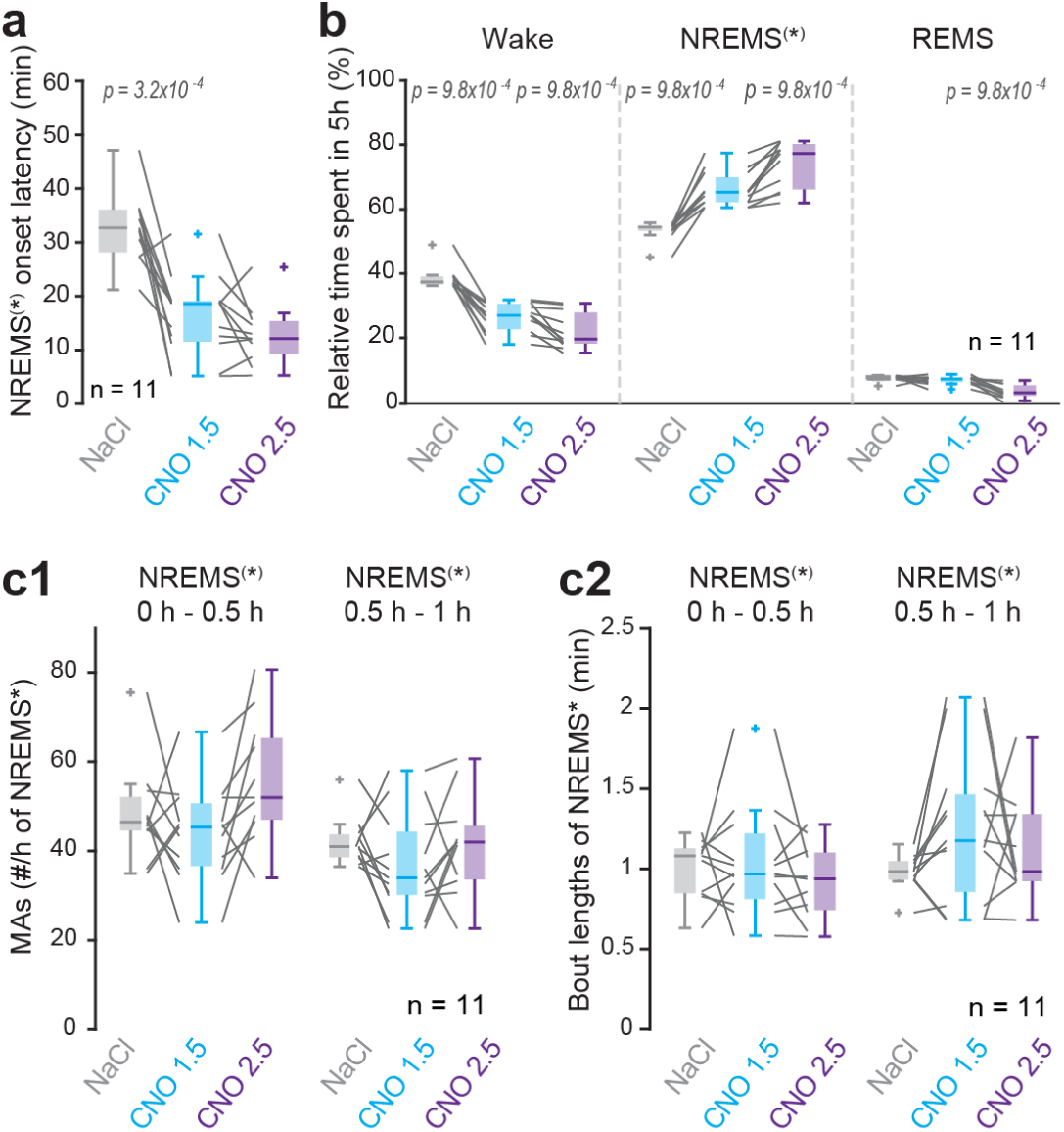
Chemogenetic activation of VSNs alters the architecture of sleep. a. Box-and-whisker plot of latencies to enter NREMS or NREMS* (together symbolized as NREMS^(^*^)^ on the y-axis) after excitatory chemogenetic stimulation of VSNs in the L-JNG. For control data, see **Supplementary Figure 5**. Grey lines connect datapairs. Repeated-measures ANOVA with factor ‘Treatment’, with post hoc t-tests and Bonferroni correction (p = 0.025); significant p values are indicated. b. Same dataset as used in a, analyzed for times spent in Wake, NREMS or NREMS* and REMS during 5-h recordings starting with i.p. injections. One-way ANOVA or Friedman test with factor ‘Treatment’ separately for Wake, NREMS^(^*^)^, REMS, with *post hoc* t-test or Wilcoxon signed-rank tests and Bonferroni correction (p = 0.025), significant p values are indicated. c. As a, for microarousal (MA) density (c1) and bout lengths (c2). Data analysis was done in 30-min bins, which allowed to compare equal NREMS or NREMS* times for data from NaCl and CNO injections. One-way repeated-measures ANOVA or Friedman tests with factor ‘treatment’ for c1 and c2, with Bonferroni correction (p = 0.013). No significant p value.

### Chemogenetic stimulation of VSNs preserves homeostatic regulation of NREMS* and REMS

Both NREMS and REMS are regulated homeostatically, such that changes in the time spent in each of the two states drives state-specific compensatory modifications in the duration and/or the intensity of that state (Franken, 2002; Franken & Dijk, 2024; McCarthy et al., 2016). To test whether the homeostatic response to time spent in NREMS* differs from that of NREMS, we extended EEG/EMG recordings for up to 48 h (2 consecutive light-dark periods) after CNO injection (n = 9). We used the higher CNO dose (2.5 mg/kg) to strongly increase NREMS*, with the goal to optimally detect changes in sleep times during the recording. The time spent in NREMS* or NREMS was higher after the CNO compared to the NaCl injections in the 1^st^ light phase and in the 1^st^ and 2^nd^ dark phases (**Figure 6a**, **Supplementary Figure 6a**, n = 9). We next analyzed spectral power dynamics in the δ (1.5 – 4 Hz) range during NREMS - an established measure of sleep need that dissipates exponentially with time spent in NREMS while increasing with wake (Franken & Dijk, 2024). In CNO compared to NaCl conditions, δ power decayed more rapidly in the 1^st^ light phase and mean levels remained lower throughout the 1^st^ dark phase relative to the NaCl condition (**Figure 6b**, **Supplementary Figure 6b**, n = 9). We compared these experimental data to a predictive model of δ dynamics for normal NREMS (Franken et al., 2001), assuming that NREMS* had similar effects on δ power. Despite a larger predicted than observed dynamic range in both conditions, we found a remarkable similarity in that both, the immediate decrease in δ power below levels observed in NaCl injected mice in the 1^st^ light phase, and its continued lower levels during both dark phases, were reproduced (**Figure 6c**). This suggests that NREMS* drives δ power similarly to NREMS, despite the physiological differences between the two.

**Figure 6.**
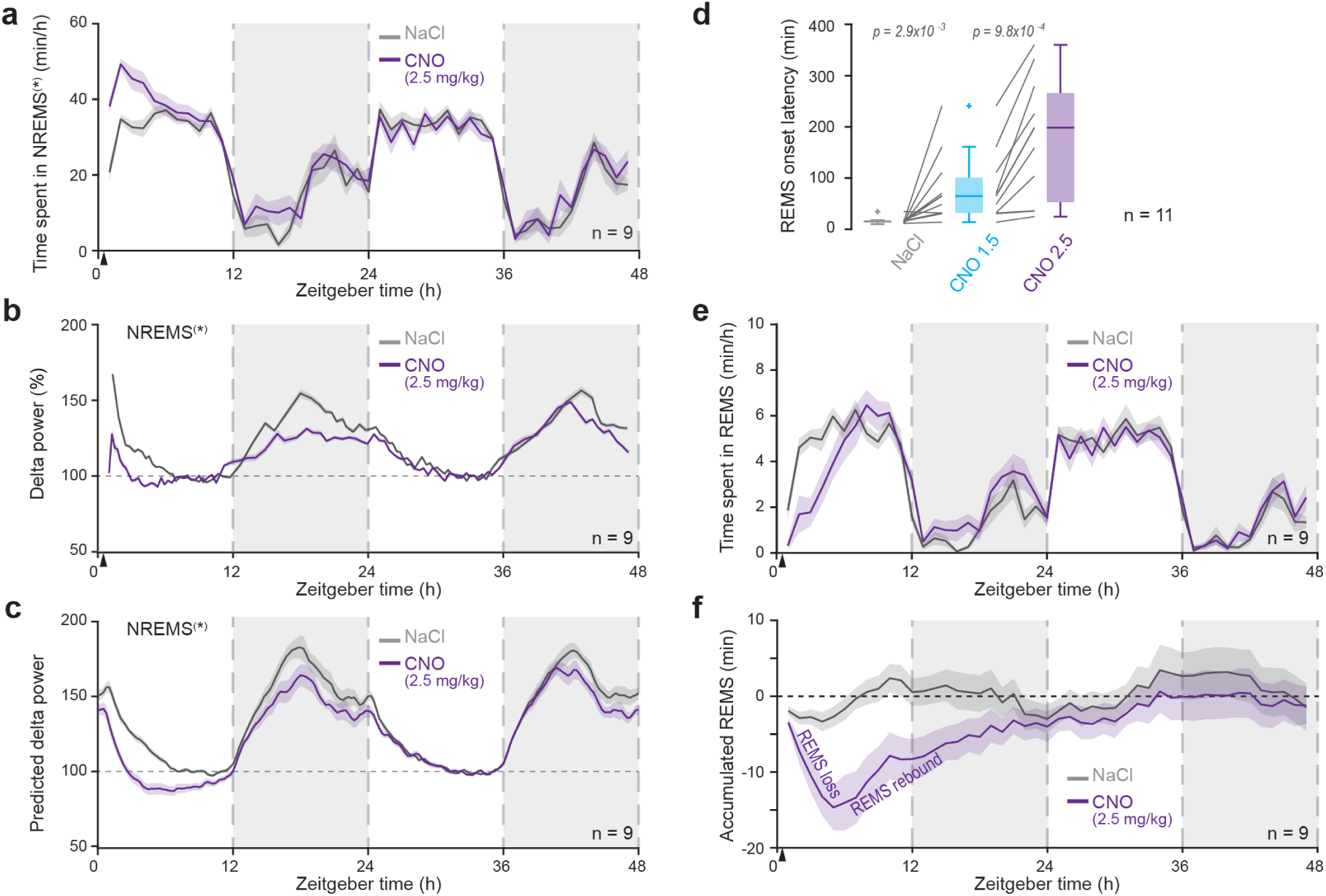
Chemogenetic activation of VSNs preserves homeostatic regulation of REMS. a. Times spent in NREMS* or NREMS (together symbolized as NREMS^(^*^)^ on the y-axis) over 48 h after NaCl or CNO (2.5 mg/kg) injections. Arrowhead denotes the time point of injection. Note increased times spent for up to 24 h (in the first dark period). See **Supplementary Figure 6a** for statistical analysis. b. Dynamics of δ power over the same period as a, calculated for 30-min bins of times spent in NREMS* and graphed in % of power values for NaCl injections for the time bins between hours 11 – 12 (end of the light phase). Arrowhead denotes the time point of injection. See **Supplementary Figure 6b** for statistical analysis. c. Simulation of δ power dynamics based on the assumption that NREMS* induces the same homeostatic process as NREMS, using published parameter estimates (Franken et al., 2006). d. Box-and-whisker plots for REMS onset latencies, calculated from the onset of NREMS for NaCl injections, and of NREMS* for CNO injections. Friedman tests for factor ‘Treatment’ with *post hoc* Wilcoxon signed-rank tests. Bonferroni corrected p = 0.025. Control data are shown in **Supplementary Figure 6c**. e. Times spent in REMS, plotted in hourly values for two light and dark phases after i.p. injections. A small loss of REMS at the beginning of the resting phase for NaCl was due to the arousing effect of the injection. Arrowhead denotes the time point of injection. f. Same data as panel e, shown as cumulative differences and calculated relative to previous 48 h-baseline recordings (without any injection). Arrowhead denotes the time point of injection. This plot emphasizes the ‘REMS loss’ that follows CNO injection but that is followed by a ‘REMS rebound’ during which lost REMS is recovered. See **Supplementary Figure 6d** for statistical analysis.

In addition to NREMS* induction, injections of CNO delayed REMS onset, with a clear dose dependence, although effect sizes varied substantially between animals (**Figures 3c, 4a**, **Figure 6d**, n = 11, see Methods), with the effect absent in control animals (**Supplementary Figure 6c**, n = 6). When tracking time spent in REMS across 48 h, we found that the loss of REMS produced by CNO was fully compensated for, with total time spent in REMS recovering to normal levels within 24 h (**Figure 6e**, **f**, **Supplementary Figure 6d**, n = 9). Monitoring of heart rate over 48 h showed that values returned to control within 12 h (**Supplementary Figure 6e**, **f**, n = 9). Therefore, the REMS suppression induced by activation of VSNs was homeostatically compensated for when CNO was cleared from the body.

### VSN stimulation lowers cortical and core body temperatures

Large ambient and/or internal temperature drops reduce EEG amplitude (Harding et al., 2018; Krauchi & Deboer, 2010; Kroeger et al., 2018) and suppress REMS (Amici et al., 2008; Harding et al., 2018; Komagata et al., 2019; Kroeger et al., 2018). To investigate whether the VSN stimulation lead to a temperature change, we measured cortical and body temperatures together with EEG/EMG in a separate group of Vglut2-Cre animals expressing excitatory Dreadd-receptors in L-JNG neurons (**Figure 7a**, n = 7). Baseline recordings confirmed that cortical temperature variations with sleep and wake states were in line with previous reports (Hoekstra et al., 2019), declining during NREMS and transiently increasing with every REMS bout (**Figure 7b**-**c**). Compared to NaCl, VSN stimulation by CNO produced a drop in cortical temperature that was substantially longer and larger (**Figure 7d**, **e**, n = 7).

**Figure 7.**
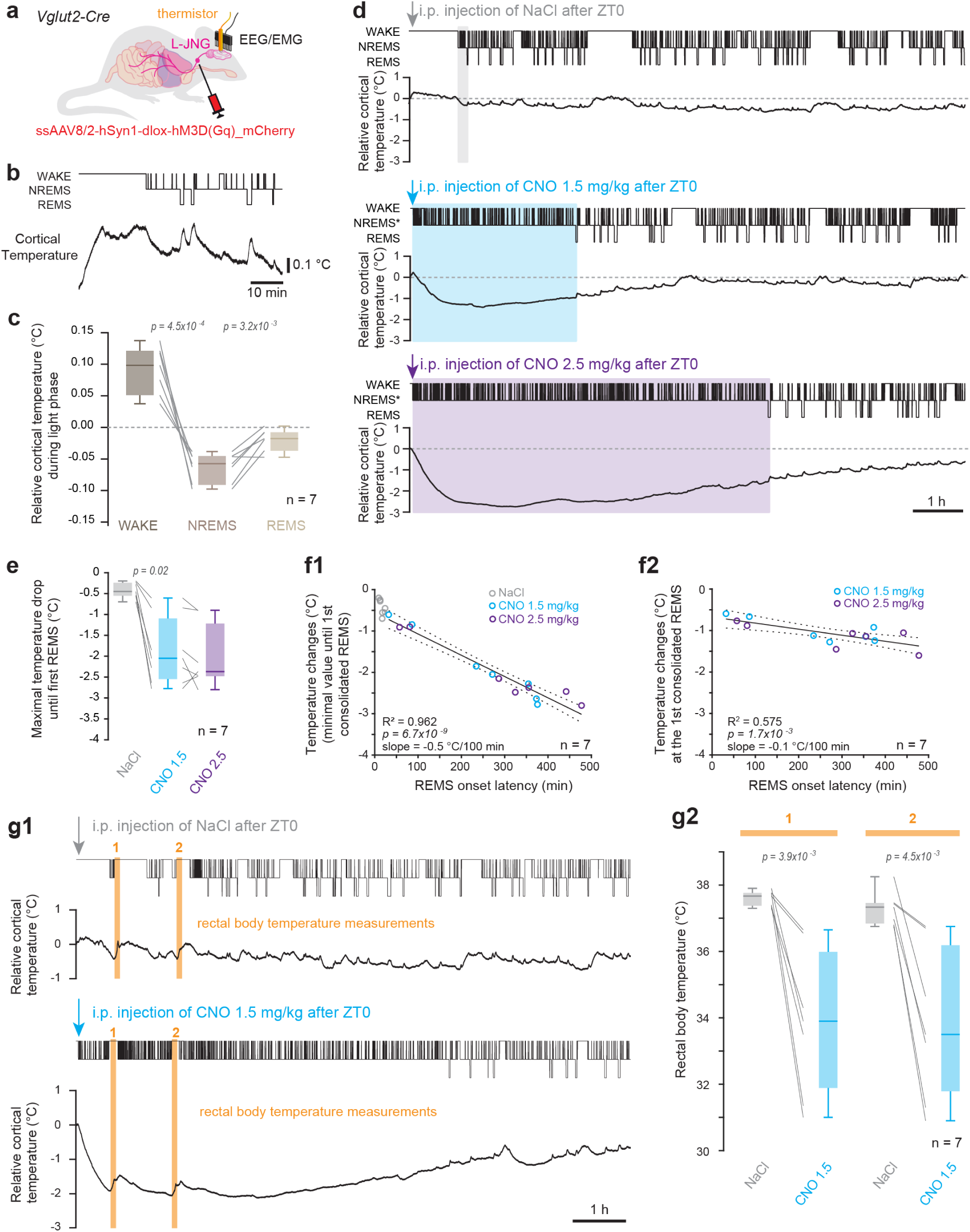
A chemogenetic VSN stimulation induces transient brain-body temperature decrease. a. Schematic of the viral injection and surgeries for combined VSN stimulation, EEG/EMG recordings, and cortical temperature measures. b. Example recording of cortical temperature during spontaneous sleep-wake behavior before VSN stimulation. c. Box-and-whisker plots of mean cortical temperature values for wake, NREMS and REMS, calculated relative to the mean temperature values for each animal across the three states during the light phase. One-way ANOVA with factor ‘Vigilance state’, followed by post hoc t-tests, Bonferroni-corrected p = 0.025. Significant p values are indicated. d. Example cortical temperature recordings from one mouse. Top: Hypnogram; Bottom: cortical temperature expressed as the difference to the temperature at NREMS(*) onset. Colored rectangles show time to the first consolidated REMS period. e. Box-and-whisker plots of maximal temperature drops between NREMS* onset and the first consolidated REMS. Friedman test with factor ‘Treatment’, with *post hoc* Wilcoxon signed-rank tests and Bonferroni corrected p = 0.025. Significant p value is indicated. f. (f1) Linear regression between temperature drops and REMS onset latency, yielding a slope = -0.5 °C/ 100 min; R^2^ = 0.962, p = 6.7 * 10^-9^. (f2) Linear regression between relative temperature at which REMS re-appears and REMS onset latency, with slope = -0.1 °C/ 100 min; R^2^ = 0.575, p = 0.0017. g. (g1) Example cortical temperature recordings combined with rectal temperature measures at the 2 time points indicated by vertical yellow bars. Rectal measurements woke up the animals and transiently increased cortical temperature. (g2) Box-and-whisker plots for corresponding mean rectal temperature values sampled at the times shown in g1. Paired t-tests, with Bonferroni corrected p = 0.025.

The cortical temperature change correlated negatively with the delay to REMS onset for all datapoints obtained from the two CNO doses (**Figure 7f1**, n = 7), suggesting that the recovery of REMS depended on the return to normal temperature. In support of this, the temperature at which REMS re-appeared varied less with the delay to REMS onset, consistent with a threshold above which REMS could be again expressed (**Figure 7f2**, n = 7) (see Discussion). Rectal probes further showed that core body temperatures were also lowered by VSN stimulation (**Figure 7g1**, **g2**, n = 7). This finding is in line with similar core and brain temperature drops found with electrical stimulation of the entire vagus nerve in rats (Larsen et al., 2017) and with chemogenetic stimulation of a VSN subpopulation (Scott et al., 2025).

### Ambient warming identifies brain-body cooling as major effector of VSN stimulation

Given the observed brain-body cooling, we hypothesized that ambient warming could counteract the observed actions on sleep. We warmed animals through heating pads positioned within and below the cages for the first hour after CNO (1.5 mg/kg) injection (**Figure 8a**). As a result, the CNO-induced cortical temperature decline was reduced (**Figure 8b**, **c**, n = 8). Furthermore, the latency to REMS onset, albeit very strongly shifted in this group of animals, was shortened (**Figure 8d**, n = 8) and the spectral dynamics typical for NREMS recovered more quickly (**Figure 8e1**, **e2**, **Supplementary Figure 7**, n = 8). The negative correlation between cortical temperature drops and REMS onset latency was preserved in the case of heating, again supporting the temperature drop as one of the primary effectors of VSN stimulation (**Figure 8f1**). In line with this, the temperature at which REMS re-appeared remained constant across REMS onset latencies (**Figure 8f2**). Therefore, counteracting the temperature decline accelerated the recovery of the major effects of VSN stimulation on sleep.

**Figure 8.**
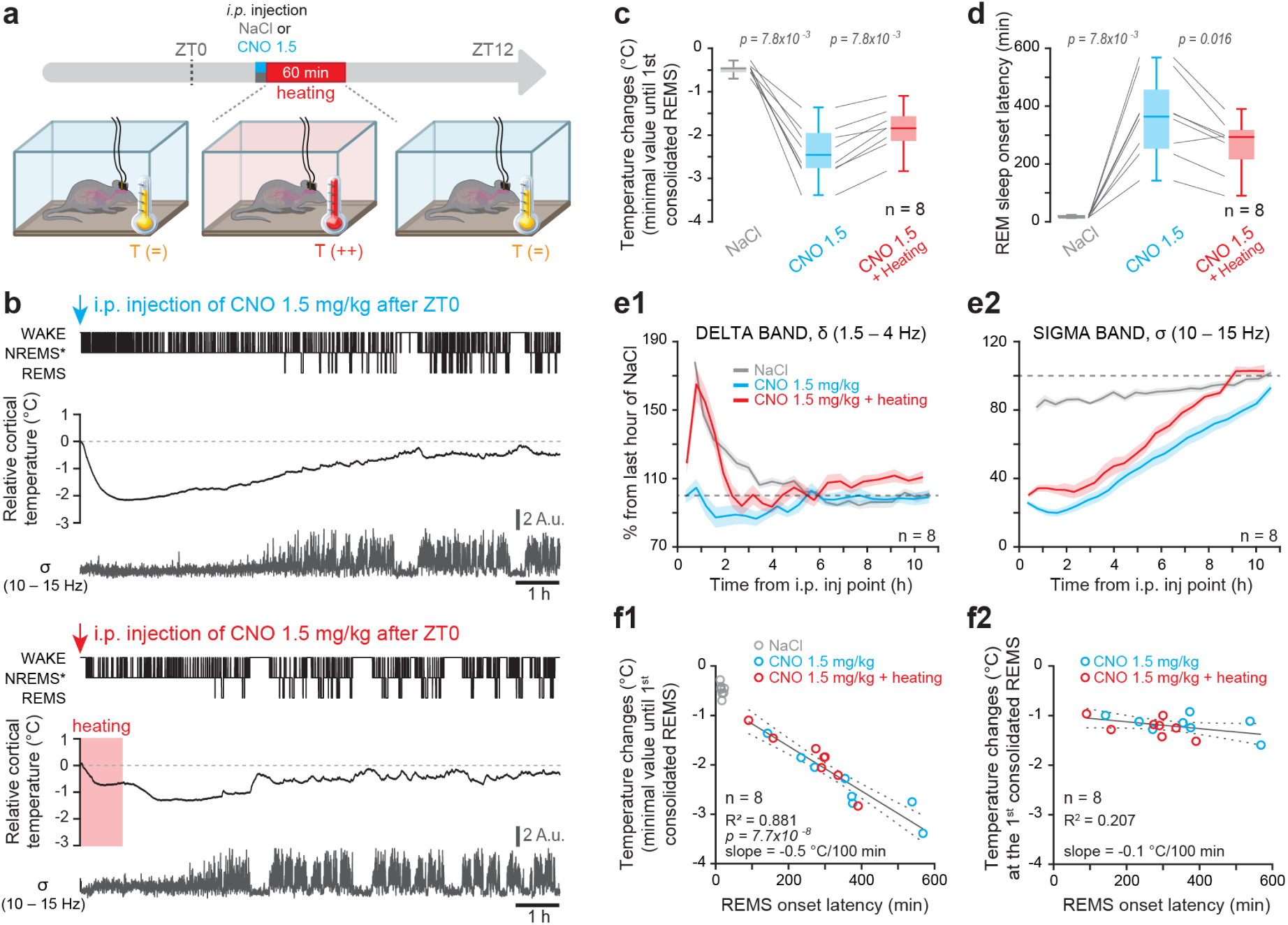
Prevention of VSN stimulation-induced brain-body cooling by ambient warming sped up recovery to normal sleep. a. Schematic of experimental set-up. For the heating procedure, see Methods. b. Example recordings from one mouse in the absence (top) or presence (bottom) of external warming (red shaded rectangle). From top to bottom, hypnogram, cortical temperature and the dynamics of σ (10 – 15 Hz) power. c. Box-and-whisker plot for cortical temperature drops across experimental treatments. Friedman test with factor ‘treatment’ (NaCl, CNO, CNO+heating), *post hoc* Wilcoxon signed-rank tests with Bonferroni-corrected p = 0.025. Significant p values are indicated. d. As c, for REMS onset latencies across experimental treatments, quantified as in Figure 6d. Statistics as in c. Note that some of the animals responded to CNO (1.5 mg/kg) with a very long REMS onset latency, nevertheless, ambient warming clearly reduced it. e. Spectral dynamics across experimental treatments, calculated as in Figure 4c. See **Supplementary Figure 7** for statistics. f. (f1) Linear regression between REMS onset latency and temperature drop, yielding a slope = -0.5 °C/ 100 min; R^2^ = 0.881, p = 7.7 * 10^-8^. (f2) Linear regression between relative temperature at which REMS re-appears and REMS onset latency, with slope = -0.1 °C/ 100 min; R^2^ = 0.207, not significant.

## Discussion

Sleep has important benefits for brain and body health. The activity of VSNs may contribute to these benefits by regulating brain-body correlates of sleep. However, a comprehensive description of these correlates has yet to be done. To this end, we collected behavioral, polysomnographic, temperature and neural activity measures in mice while stimulating VSNs during their preferred resting period. Polysomnography indicated the presence of spectral hallmarks of NREMS during VSN stimulation, together with behavioral signatures of maintained arousability. Like NREMS, we observed drops in heart rate and brain-body temperature. However, in contrast to NREMS, sleep spindles were absent and there were no transitions to REMS. In further opposition to NREMS-like characteristics, there was a marked increase in noradrenergic activity, akin to a wake-like pattern. Vagal sensory activation can thus blend features of NREMS and wakefulness, generating a state of global sedation and brain-body cooling while behavioral and neuronal signatures typical for wakefulness are present. At the same time, our data also suggest that VSN stimulation, if finely tuned, may be useful to regulate sleep along a continuum of variables to possibly enhance its health-promoting effects.

The simultaneous presence of sleep- and wake-related features supports our initial hypothesis that VSN activation may influence sleep through its actions on both the autonomic nervous and the brain’s arousal systems. Still, the opposition between LC activation in an otherwise sedated organism seems paradoxical. One aspect to consider is that the VSN stimulation-induced actions may reciprocally interact after VSN activation started. Most importantly, declines of body temperature are well-known as a protective response in small mammals in response to environmental or energetic challenge (Amici et al., 2008; Krauchi & Deboer, 2010; Lovelace et al., 2023; Scott et al., 2025). Therefore, chemogenetically elevated VSN activity may, in part, decrease temperature not only as part of its actions on the body but also because it constituted a challenge for the mice. However, body temperature decreases were observed in rodents in response to various stimulations of the vagus nerve, differing in intensity and duration (Larsen et al., 2017; Lovelace et al., 2023; Scott et al., 2025). Therefore, to what extent temperature decreases may be a generic response of small mammals to elevated body-brain communication is an important point to address, for example through diverse electrical, opto- and chemogenetic VSN stimulation protocols. In any case, our findings ask for caution when studying brain-body communication in small mammals, as hypothermia could be a confounding factor that can go overlooked.

Although we denoted it as a NREMS-like state, the VSN-induced state was different from NREMS in its physiological manifestations. First, the cooling of cortical surface and core body temperature associated with the NREMS-like state was stronger than that for NREMS (Deboer et al., 1994; Hoekstra et al., 2019). A core body temperature drop was also described after stimulation of VSN subpopulations innervating heart or the gastrointestinal system (Lovelace et al., 2023; Scott et al., 2025). This stronger temperature change calls for a comparison to states of torpor. Daily torpor of mice and hamsters refers to an adaptive state to cope with energetic challenges of fasting and environmental cooling (Ambler et al., 2021; Deboer & Tobler, 1994; Huang et al., 2021). In animals housed at room temperature as ours, torpor results in core temperature declines (∼10 °C) (Deboer et al., 1994; Hrvatin et al., 2020) that are larger than what we observe (∼4°C) with VSN stimulation. It remains to be seen whether the NREMS-like state differs from torpor in the factors leading to a temperature decrease. The stimulation of VSNs as done in our study plausibly leads to a strong parasympathetic activation that increases blood flow to the periphery and dilates distal vessels (Krauchi & Deboer, 2010; Szymusiak, 2018). The resulting heat loss may overwhelm the normal cold defense and lower core body temperature. The decrease of cardiovascular activity could further prompt cooling as part of an energy-saving reaction (Scott et al., 2025). Additionally, hypothermia can be induced via AP neurons (Engström Ruud et al., 2024), which we found activated after VSN stimulation. Therefore, while the phenomenological descriptions of ‘torpor-like’ or ‘NREMS-like’ may both apply to the VSN-stimulation-induced state, more work is needed to establish its relation to torpor and to other thermoregulatory mechanisms (Hrvatin et al., 2020; Krauchi & Deboer, 2010). Conversely, it would be interesting to determine whether VSN activity is a possible contributor to hypometabolism-induced temperature declines in torpor states.

In a second noteworthy contrast to NREMS, we observed elevated cFos activity in the LC and increased population activity in noradrenergic neurons. Previous studies using vagus nerve stimulation indicated higher noradrenergic activity as part of the response (Berger et al., 2021), which is commonly thought to involve activation of the LC via ascending circuits from NTS (Van Bockstaele et al., 1999). However, the combined data from this study suggest that an alternative scenario may need to be considered. LC activation could be, in part, result secondarily from VSN-induced effects on heart rate and temperature. This is supported by the observation that LC activity and temperature declines recovered in parallel with the sleep architectural changes. Additionally, LC activity has been shown to increase in response to internal state changes, such as declines in blood pressure (Valentino et al., 1991), which could facilitate arousal (McGinley et al., 2015; Page & Valentino, 1994) and promote vasoconstriction (Hauglund et al., 2025) to counteract the decrease of cortical temperature. LC neuronal excitability appears to be only weakly sensitive to brain cooling (Corrodi et al., 1967). These findings suggest that LC activity could act as a central alarm response to the temperature and cardiovascular effects provoked by VSN stimulation, in an effort to sustain arousability. Such a constellation of factors leading to LC activation remains to be explored. Yet it is noteworthy that animals undergoing stimulation of only a subgroup of VSN neurons show heightened anxiety levels, which could reflect LC activation (Scott et al., 2025). As LC is also likely activated by vagus nerve stimulation in humans (Berger et al., 2021), a better understanding of the causes and consequences of such activation will be relevant for further treatment options.

In a third contrast to NREMS, the VSN-induced NREMS-like state did not show transitions to REMS. In line with the temperature sensitivity of REMS (Amici et al., 2008; Harding et al., 2018; Komagata et al., 2019; Kroeger et al., 2018), we found a tight correlation of REMS onset with the temperature decline across experimental conditions without or with ambient heating. This is a strong indication for a dominant role of temperature in delaying REMS during VSN stimulation. Even more so, the temperature at which REMS re-appeared varied minimally, consistent with a threshold temperature above which REMS is again possible. This threshold temperature became more negative for long REMS latencies, probably because the need to enter REMS was then very high. Aside temperature, the persistent activation of LC could have contributed to REMS suppression, which remains to be tested.

In spite of the pronounced physiological differences with NREMS listed above, during the NREMS-like state, the homeostatic regulation appeared unaltered. Thus, the initial drop of low-frequency (δ) power after VSN stimulation could be explained by the extra time spent in the NREMS-like state without a need to infer temperature and/or CNO effects on the expression of δ power nor on its relation with prior sleep-wake history. This argues against a major contribution of additional cooling/CNO effects on the euthermic sleep-wake driven dynamics of δ power and contrasts with a slow-down of sleep homeostasis in torpor (Heller & Ruby, 2004; Krauchi & Deboer, 2010; Strijkstra & Daan, 1997) or dexmedetomidine-induced cooling (Ma et al., 2019) and supports the use of ‘NREMS-like’. A further noteworthy observation is the persistence of enhanced NREMS time long after CNO has been cleared. This illustrates the proposed separate regulation of δ power and NREMS time in the homeostatic response (Franken & Dijk, 2024). Moreover, the homeostatic regulation of time spent in REMS was not suspended during the NREMS-like state and the loss of REMS was completely recovered 24h after CNO administration. Interestingly, REMS homeostasis was also preserved during a body temperature drop (and associated REMS loss) induced by cold exposure (Amici et al., 2008) or direct chemogenetic stimulation of preoptic hypothalamic galaninergic neurons (Kroeger et al., 2018). Therefore, while the temperature sensitivity of REMS expression might be adaptive (Blumberg et al., 2020; Harding et al., 2020), its homeostatic regulation seems not to be affected by brain temperature. These insights will inform the study of the circuits underlying REMS and its homeostatic regulation, for which some are shared (Maurer et al., 2024).

This study highlights that VSN activity can powerfully modulate sleep when stimulated chemogenetically. We acknowledge that this approach has led to an intense stimulation of VSNs that presented vital challenges for the animals. The results bring to light that we know little about the natural activity patterns of VSNs, with novel technologies to measure it in development (Marmerstein et al., 2022) and evidence that this activity varies across the light-dark cycle just starting to be recognized (Borgmann et al., 2021). We further identify important differences but also resemblances of the NREMS-like state to natural sleep showing that multimodal assessment of brain-body physiology is necessary to identify the key driving factors. It is likely that VSNs from different organs might affect sleep in specific ways that we can now start dissecting taking advantage of the genetic tractability of VSN neurons (Prescott & Liberles, 2022). Future work will tell whether this may unfold a map of sleep profiles that may inform about bodily health and inspire approaches to sleep diagnosis and treatment in humans. Moreover, as novel medications for weight management such as semaglutide may act on vagal activity, our work provides a timely contribution to consider potential side effects of these on sleep.

## Acknowledgements

We are grateful to Myrtha Arnold and Jean-François Krieger (ETHZ) for introducing us into vagal surgeries, and to Prof. Liberles (Harvard University) for initial technical advice. We are indebted to the numerous colleagues who provided us with insightful comments and who shared knowledge on various aspects of this study, notably Dr. J. Buron, Proff. Henning Fenselau, V. Mansuy-Aubert, C. Menuet, M. Schmidt, V. Vyazovskiy, P. Fuller and E. Arrigoni. We thank Dr. A. Kolaxi and M. Pertin for help with immunostaining protocols and Dr. L. Restivo for help in setting up the behavioral monitoring. Dr. Gil Vantomme performed the first *in vitro* recordings, and Dr. Romain Cardis provided advice for Matlab coding and statistical analysis. The Master students D. Gauthier and N. Eliasson are acknowledged for their contributions to unpublished portions of this work. The constructive input of all members of the Luthi lab is highly appreciated.

## Author contributions

NC carried out all experiments and analyses except for Figure 3, GF contributed data and analysis for Figure 3, LF contributed to temperature recordings and the analysis of cFOS expression. YE helped with scoring. YE and PF helped to set up pilot temperature recordings, AOF and LF helped with analysis and figure preparations. AL and PF conceived the original project, NC and AL wrote the first draft of the manuscript and received input from all authors.

## Material and Methods

### Animal husbandry

Mice from the Slc17a6^tm2(cre)Lowl^/J (JAX Stock#016963), commonly referred to as Vglut2-Cre line, were bred on a C57BL/6J background (breeders were kindly provided by Dr. A. Carleton, UNIGE). Mice were housed in a humidity- and temperature-controlled (22 °C) animal facility with a 12 h / 12 h light-dark cycle. For viral injections, 4 – 8 week-old animals of either sex were transferred into a ventilated cabinet in the P2 safety level facility with similar conditions on the day prior to injection. They remained in the P2 facility for 72 h after the viral injection. For *in vitro* experiments, animals were transferred into a housing room with a 12 h / 12 h light-dark cycle (lights onset at 9:00 am, corresponding to ZT0) where they remained for ≥ 3 weeks until *in vitro* electrophysiology. For *in vivo* experiments, animals remained in the P2 room for ≥ 2 weeks before surgery for sleep recordings. Once animals had received surgical implants for sleep recordings, they were housed singly in home cages equipped with tall Plexiglas walls (∼30 cm) without roof in a sleep recording room for ≥ 1 week prior to habituation and recording. Through all experimental procedures, food and water were given *ad libitum*. All experimental procedures were carried out according to the Swiss National Guidelines on Animal experimentation and are part of a license approved by the Cantonal Veterinary Office for Animal Experimentation.

### Surgery for viral injections in the L-JNG

Prior to surgery, mice were given analgesic (carprofen, 5mg/kg s.c.), anesthetized by isoflurane inhalation (5 % in oxygen), then placed on a warming surface for maintenance of body temperature at 37 °C. Anesthesia was maintained via a nose cone through which isoflurane was provided (1.5 – 2.5 % in oxygen), while the mouse was placed on its back. Some of the following surgical steps are visually illustrated in **Supplementary Figure 1**. The skin was shaved and local anesthetics (lidocaine (6 mg/kg) + bupivacaine (2.5 mg/kg, 20 µl)) injected s.c. at the location of the future incision. A 3-4 mm-long incision was made on the ventral surface of the neck via fine scissors, the salivary glands were exposed and separated. A magnetic fixator retraction system (FST, retractors with 1-mm tip diameter) was used to gently displace the different muscles lying on top of the JNGs. To expose the L-JNG, one retractor each was placed on the left salivary gland and sternomastoid muscles. A third retractor was used to displace the left sternohyoid and the omohyoid muscles to the right. Finally, a fourth retractor was placed on the left digastric muscle. The left hypoglossal nerve was gently separated from the vagus nerve through slightly lifting with a fifth retractor (0.5 mm tip diameter), which exposed the L-JNG. A manual manipulator was used to insert a thin glass pipette (5-000-1001-X, 649 Drummond Scientific) pulled on a vertical puller (Narishige PP-830), initially filled with mineral oil, and backfilled with the virus just prior to injection, into the L-JNG. The injected volume of virus ranged between 250 – 350 nL. The following injections were done into VSNs of the L-JNG in Vglut2-Cre animals:

**-** expression of ChR2: 11 animals were transfected with AAV1-EF1α-DIO-hChR2(H134R)-eYFP-WPRE-hGH (1.9×10^13^ GC/ml, Addgene) and 1 animal with AAV1-CaMKIIα-hChR2(H134R)-eYFP-WPRE-hGH (1.1×10^13^ GC/ml, Addgene)
**-** expression of excitatory chemogenetic Dreadd receptors: 21 animals were transfected with ssAAV8/2-hSyn1-dlox-hM3D(Gq)_mCherry-dlox-WPRE-hGH(pA) (×10^12^ GC/ml, VVF Zürich)
**-** expression of Dreadd-unrelated constructs to control for CNO effects: 1 animal with AAV1-EF1α-DIO-hChR2(H134R)_EYFP-WPRE-hGH (titer 1.1×10^13^ GC/ml, Addgene), 1 animal with AAV8-hSyn-FLEX-Jaws_KGC_GFP-ER2 (titer 3.2×10^12^ GC/ml, UNC Vectore Core), and 4 animals with ssAAV8/2-hSyn1-dlox-mCherry(rev)-dlox-WPRE-hGHp(A) (titer 9×10^12^, VVF Zürich)
**-** expression of jGCaMP8s in combination with chemogenetic stimulation of VSNs in L-JNG: we used the dopamine-β-hydroxylase-(i)Cre line (kindly provided by Prof. T. Patriarchi, UNIZH): 6 animals were injected with AAV5-hSyn1-dlox-jGCaMP8s (titer 5.8×10^12^ GC/ml, VVF Zürich) in the LC and with ssAAV8/2-hSyn1-hM3D(Gq)_mCherry (titer 3.0×10^12^ GC/ml, VVF Zürich) in the L-JNG. One of the animals was excluded due to poor expression of the virus injected in the L-JNG.

### *In vitro* electrophysiological recordings

#### Brain slice preparation

Coronal brainstem slices containing the NTS and AP were prepared from Vglut2-Cre mice previously injected with a ChR2-expressing viral vector as described above. 4 – 7 weeks after viral injection, mice aged 8 – 12 weeks were subjected to isoflurane anaesthesia (4 % in O2) after which they were decapitated, brains extracted and quickly immersed in ice-cold oxygenated solution with partial substitution of NaCl containing (in mM): NaCl 66, KCl 2.5, NaH2PO4 1.25, NaHCO3 26, D-saccharose 105, D-glucose 27, L(+)-ascorbic acid 1.7, CaCl2 0.5 and MgCl2 7, using a sliding vibratome (Histocom). Brains were cut at the level of bregma in the coronal plane and the posterior portion containing the brainstem was glued with the trimmed surface on an ice-cold metal blade, with the ventral side apposed to a supporting agar block. Acute 300-µm-thick coronal brainstem slices were prepared in the same ice-cold oxygenated sucrose solution, transferred to a storage chamber and kept for 30 min in a recovery solution at 35°C (in mM): NaCl 131, KCl 2.5, NaH2PO4 1.25, NaHCO3 26, D-glucose 20, L(+)-ascorbic acid 1.7, CaCl2 2, MgCl2 1.2, *myo*-inositol 3, pyruvate 2, after which they were kept at room temperature for at least 1 h. All recordings were done at room temperature.

#### Patch-clamp recordings

Recording pipettes were pulled from borosilicate glass (TW150F-4) (WPI) with a DMZ horizontal puller (Zeitz Instr.) to a final resistance of 4.6 – 6.1 MΩ. Pipettes were filled with a K^+^-based intracellular solution that contained (in mM): KGluconate 140, HEPES 10, KCl 10, EGTA 0.1, phosphocreatine 10, Mg-ATP 4, Na-GTP 0.4, supplemented with neurobiotin (2mg/mL), pH 7.3, 290 – 305 mOsm. Slices were placed in the recording chamber of an upright microscope (Olympus BX50WI) and continuously superfused with oxygenated ACSF containing (in mM): NaCl 131, KCl 2.5, NaH2PO4 1.25, NaHCO3 26, D-glucose 20, L(+)-ascorbic acid 1.7, CaCl2 2 and MgCl2 1.2.

The area containing the left NTS and AP was identified through a 10X immersion objective in transillumination microscopy, then, cell bodies were visualized through a 40X immersion objective in differential interference contrast optics. Pipette offset was zeroed just prior to establishing a membrane seal and cells patched in the whole-cell recording configuration and tested for responsiveness to optogenetic stimulation. The area permitting successful patching of cells responding to light stimulation was identified within the medial portion of the NTS adjacent to the AP (Br -7.64 to -7.32 mm). In this area, immunoenhancement of EYFP also revealed a high density of green fibers, indicating that the ChR2_EYFP protein was expressed up to the terminals of the sensory axons (**Figure 1d2**, **f1**).

Signals were amplified using a Multiclamp 700B amplifier, digitized via a Digidata1322A and sampled at 10 kHz with Clampex10.2 (Molecular Devices). Passive cellular properties and action potential discharge frequencies were measured using direct somatic current injections in the voltage- or the current-clamp recording configurations. Cells included for analysis had access resistances between 21 and 48 MΩ, which were not compensated for. A liquid junction potential of -10 mV was also not compensated for.

#### Recording protocols

After gaining whole-cell access, cells were first held in voltage-clamp at -60 mV and hyperpolarized in brief steps (10 mV, 100 ms) after which whole-field blue LED (Cairn Res) light pulses (470 nm, duration: 1 – 5 ms, maximal light intensity 3.8 mW, 0.75 mW/mm^2^) were given to check for a synaptic response. Light pulses were given maximally once every 5 – 10 s for 5 – 10 sweeps. Once synaptic connectivity was established, cellular action potential discharge properties were tested in current-clamp mode using somatic current injections from -50 to +80 pA in square current steps lasting 0.5 – 2 s. Evoked responses were then again measured in voltage-clamp at a frequency of 0.1 – 0.2 Hz. The effects of DNQX (10 µM) were tested on EPSPs at 0.1 – 0.2 Hz in current-clamp mode at membrane potentials of -68 to -60 mV. Per slice, only one synaptically connected cell was studied. At the end of the recording, sections were fixed in PFA 4% in PBS for *post hoc* recovery of the neurobiotin-filled cells (Vector Labs) and for immunoenhancement of EYFP-expressing fibers.

#### Analysis of patch-clamp data

Passive cellular properties including membrane time constant (τm), cellular capacitance (Cm) and resting membrane potential (RMP) were analyzed according to standard procedures (Vantomme et al., 2020). The membrane resistance was not directly calculated from the steady-state current response due to time-dependent current components appearing even with small hyperpolarizations from a holding potential of -60 mV (Miles, 1986), and due to very high levels of spontaneous synaptic currents. However, estimations from τm and Cm indicate that these resistances were high (> 600 MΩ), as also reported previously (Kline et al., 2002). Action potential discharge was quantified by measuring action potential amplitude from the inflection point to the peak and by measuring the mean frequency of discharge in suprathreshold current injections lasting 2 s and quantified as a function of current amplitude. All neurons showed tonic action potential firing with minor discharge adaptation. EPSCs were quantified in terms of latency to test for the presence of putative mono- or polysynaptic responses (Doyle & Andresen, 2001). The latency was characterized for its jitter that was calculated as the standard deviation of six sequentially evoked responses. In most cases, EPSCs were composed of multipeak events, accordingly only the amplitude of the first peak response was measured. DNQX was applied after a stable baseline of 12 consecutive evoked EPSPs at inter-stimulus intervals of 5 s. The possible presence of inhibitory synaptic currents was not tested. The *in vitro* data were manually analyzed using Clampfit v10.2 cursor measurements and monoexponential fitting procedures.

### Histological procedures

#### Immunohistochemical staining of extracted ganglia

Access to the L-JNG was obtained via the surgical procedure described above. The L-JNG was extracted by cutting the vagus nerve with fine forceps, placed in PFA (4 %) for 60 – 90 min, and then washed in phosphate buffer (PB, 0.1 M) before incubation in sucrose (30 % in PB) overnight. The L-JNG was put in Tissue-Tek® and frozen at -80 °C to be cut on a cryostat at a thickness of 12 µm. Immunofluorescence was done on the sections using as first antibody anti-VGLUT2 raised in mouse (Merck Millipore, MAB5504) and as second antibody Cy3 goat anti-mouse (Jackson, 115-165-003) (**Figure 1a**).

#### Immunohistochemical staining and fluorescent microscopy of vagal sensory fibers and brainstem neurons

##### Recovery of cellular morphology of patch-clamped neurons and immunoenhancement of vagal sensory fiber-expressed EYFP (**Figure 1d2**, **f1**)

The PFA-fixed 300 µm-thick brain sections were first washed in PBS for 30 min, then washed in PBS-containing Triton 1% for 30 min. After being incubated in blocking solution (PBS-containing Triton 1 % and normal goat serum 2 %) for 30 min, slices were incubated for 5 days with the primary antibody (rabbit anti-EYFP, 1:3200, Lucerna Chem, STJ97104) at 4 °C for immunoenhancement of vagal sensory fibers. The sections were washed in PBS and incubated in the secondary antibody (goat anti-rabbit Alexa488, 1:500, Jackson ImmunoResearch, 111-545-003) and in Streptavidin-coupled Alexa594 (1:8000, Lucerna Chem, STJ16100613) diluted in PBS-containing Triton 0.3 % and normal goat serum 2 %, for 1 day at 4°C. Before mounting, the sections were washed in PBS for 45 min.

##### cFos protein staining after chemogenetic stimulation of VSNs (**Figure 2c1**, **3a1**, **Supplementary Figure 2**)

Animals were i.p. injected with NaCl or CNO (1.5 mg/kg) around ZT0, brains extracted after 60 – 90 min, perfused with 4% PFA for 1 day, and cryoprotected in 30% sucrose-containing PB before being cut in a cryotome at 50-µm thickness. Sections were first washed in PBS during 30 min and then in PBS-containing Triton 0.3 % for 30 min before being incubated in a blocking solution of PBS-containing Triton 0.3 % and normal goat serum 2 % for 1 h. After overnight incubation in the primary antibody (rabbit anti-cFos, 1:1000, BioConcept, 2250S) at 4 °C, the sections were washed in PBS-containing Triton 0.3 % for 30 min and then incubated in the secondary antibody (goat anti-rabbit Alexa488, 1:300, Jackson ImmunoResearch, 111-545-003; or donkey anti-rabbit Alexa647, 1:300, Thermo Fisher Scientific A-31573) for 90 min. Then, they were washed in PBS for 20 min, incubated in PBS-containing Hoechst 1 % for 10 min for nuclear staining, then washed again for 10 min and mounted. A confocal microscope (Leica Stellaris 8) equipped with a 63x oil objective (HC PL APO 63x/1.40 oil CS2) was used to acquire the images through red, green, and blue emission channels. Detection and counting of the fraction of cFos-expressing neurons over DAPI-detected nuclei was done using a machine learning-based approach in QuPath (version 0.5.1). The random forest classifier was trained on images from NaCl and CNO 1.5 mg/kg injection.

### Behavioral monitoring

Mice were video tracked (Ethovision 17.0 XT, Noldus) in their own cage and the total distance moved per minute was calculated during 150 min, starting 20 min after NaCl or CNO (1.5 mg/kg) injection (**Figure 2g**, **h**). For the loss-of-righting reflex, animals were gently lifted from their cage at 60 min after injection and placed on their back inside a V-shaped piece of wood. Animals were observed for 30 s and scored as negative for the loss-of-righting-reflex when they turned themselves back on their paws (Gelegen et al., 2018).

### Thermistor calibration

Thermistors (Digikey thermistors (SC30F103W, 10 kΩ, epoxy sensor in polyimide sleeve) were connected to a custom-made constant current source of 100 µA and immersed into a water bath held at either 25 °C or 37 °C. Once the thermistor was inserted in the water and the temperature was stable, the heating of the water bath was turned off to avoid vibrations and voltage measures were then taken immediately for 20 s. Current sources were tested before for stability over a time period of 12 h. Voltages across the thermistor were measured for the two different temperatures and the corresponding resistances (R) calculated using Ohm’s law. For every thermistor, a material constant was calculated according to the equation with temperature values at 25 and 37 °C in °Kelvin (T25= 25°C + 273.15°, same for T37), following the procedure by (Hoekstra et al., 2019):

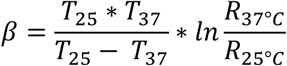

The cortical temperature (t) in °C could then be calculated according to the equation, with Rt the resistance at t measured experimentally:

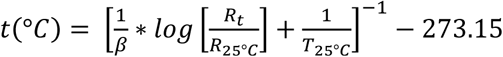

### EEG/EMG Surgeries combined with thermistor implantation

#### Surgeries for EEG/EMG recordings (**Figure 2f, 3-6**, **Supplementary Figures 3-7**)

The surgeries for implantation of EEG/EMG electrodes were done ≥ 2 weeks after the viral injection in the L-JNG. After pre-treatment with Carprofen (5 mg/kg s.c.) and induction of isoflurane anesthesia (5 % in oxygen), animals were fixed on a stereotaxic apparatus (Kopf). Anesthesia was maintained at 1.5 – 2 % of isoflurane in a mixture of O2 and N2O. An incision was made to expose the skull and the bone was scratched with a scalpel blade for a good adhesion of the head implant. After small craniotomies (0.3 – 0.5 mm) over the left frontal and parietal regions, two gold-coated copper-wire electrodes were placed on top of the *dura mater* that serves as EEG electrodes. A silver wire (Harvard Apparatus) was inserted into the occipital bone over the cerebellum for neutral reference (without touching the dura mater). For EMG recordings, two gold wires were inserted into the neck muscles. All electrodes were glued (Loctite Schnellkleber 401) and soldered to a multisite connector (Barrettes Connectors 1.27 mm, male connectors, Conrad).

#### Surgeries for combined EEG/EMG and cortical temperature recordings (**Figure 7, 8**)

We combined EEG/EMG surgery with implantation of miniature thermistors. Thermistors were placed on the cortical surface via a craniotomy drilled at coordinates (relative to Br in mm): AP -2.5, L -2.5 on the left hemisphere. To minimally damage the tissue, the thermistor was inserted by maximally 1 mm, leaving 5 mm outside. The thermistor was then glued to the skull and embedded in dental cement together with the EEG and EMG electrodes, letting the two wires protrude for later direct connection to the current generator.

#### Surgeries for EEG/EMG recordings combined with fiber photometry (**Figure 3**)

For monitoring LC activity, DBH-(i)Cre mice previously injected with excitatory Dreadd receptor-expressing virus in the L-JNG underwent a surgery for implantation of EEG/EMG recordings as described above and an optic fiber over the right LC similar to the one described in (Osorio-Forero et al., 2024). The AAV5-hSyn1-dlox-jGCaMP8s virus was injected in the right LC at 5 different depths (coordinates are given relative to the Br in millimeters lateral (L), 0.95; anterior–posterior (AP), −5.4; dorsal–ventral (DV), -4.0 to –3.2 with a step of 0.2 mm, 100 nl per depth, 500 nl in total). The optic fiber was then positioned over the injection area at a depth of -3.2 mm. After the fiber implantation, a set of EEG/EMG electrodes was implanted as previously described.

#### Timeline of the combined chemogenetics and sleep recordings experiment

Once ready for recording, mice were first recorded for 48 h baseline recordings without any injections. Mice were then injected in groups of 4 by either NaCl, CNO (1.5 mg/kg) or CNO (2.5 mg/kg) in alternating recording sessions, with the experimenter blinded to the injection solution. Every recording started at ZT0 with injections taking place within the first 45 min. The mice were then left undisturbed in their home cage and recorded for at least 5 up to 48 h depending on the experimental series (**Figure 2f, 3-8**, **Supplementary Figures 3-7**). Per CNO dose, at least 1 – 3 recordings were done per mouse, NaCl injections were carried out 2 – 4 times interspersed between the CNO conditions. All animals included for sleep effects in response to chemogenetic stimulation were perfused at the end of the experiment and underwent a fluorescent microscopic verification for the presence of vagal sensory fibers in the NTS as described above. In case no fibers could be detected, the animals were excluded from the experiment. Despite consistent actions of CNO on measured sleep parameters, effect sizes remained variable, as for example evident in the REMS latency (**Figure 6d**). Furthermore, when effects of CNO at 1.5 mg/kg were strong, there was little difference to the ones at 2.5 mg/kg (**Figure 7f**, **8f**). We attribute this variability to difficulties in controlling the extent of viral transfection of L-JNG neurons, in part because virus-containing liquid injected into the ganglion could leak outside the ganglion while we sutured the throat.

### EEG/EMG and cortical temperature recordings

After a recovery period of 1 week, animals were habituated to the cabling and to i.p. injections (of NaCl) for 5 – 7 days, followed by 48 h-baseline recordings (**Figure 7, 8**). All the signals (EEG frontal and parietal, left and right EMG, S1 LFP) were acquired at 1 kHz using an Intan digital RHD2132 amplifier board and a RHD2000 USB Interface board connected with SPI cables (all Intan Technologies) via a custom-made support system (Homemade adapters containing an Omnetics A79022-001 connector linked to a female Conrad Barrettes Connector). The data were acquired with MATLAB (RHD2000 MATLAB toolbox and a customized software). When we combined sleep recordings with cortical temperature measures, animals were connected to the same current source used before for thermistor calibration and 100 µA-currents were injected from the onset of the recordings, while recording the voltage signal at 1 kHz through an analog digital input of the Intan RHD2132 amplifier board.

### Rectal body temperature measures

We measure body temperature during combined sleep and cortical temperature recordings using a fine rectal probe (Physitemp RET-3 rectal probe for mice) (**Figure 7g1**, **g2**). Animals were previously habituated to the procedure for ≥ 5 days. To measure body temperature, the animal body was gently lifted by its tail and placed on its 4 paws. The tail was bent above the body to make the rectum accessible. The probe was previously covered with Vaseline© and gently inserted into the rectum to a fixed depth (2 cm). Stable temperature recordings were obtained within ∼10 s after insertion. The probe was disinfected after every use. During recordings, rectal temperature was measured at two time points, 45 min and 2 h after the i.p. injection of either NaCl or CNO. Recordings with rectal temperature measures were done in separate sessions not included in the analysis on CNO-induced effects on sleep and cortical temperature.

### Procedure for ambient warming

To prevent the hypothermia induced by vagal sensory stimulation, we used heating pads placed inside and below the cage that we turned on at the time of the i.p. injection for 1 h (**Figure 8**). The heating pad was previously tested such that it prevented the decline in rectal temperature by ∼75 % (without heating pad the body temperature drop was -3.6 °C whereas it was -0.9 °C with the heating pad, n = 2). For this effect, the heating pads had to be heated to 37 – 39 °C.

### *In vivo* data analysis

#### Scoring of vigilance states

Sleep scoring was done according to established procedures in the lab in a manner blinded to the treatment, using a custom-made software developed in MATLAB (MathWorks) available on GitHub (https://github.com/luthilab/IntanLuthiLab). The three major vigilance states Wake, NREMS and REMS were scored in 4-s epochs according to the following criteria. Wakefulness was identified based on combined high EMG activity and a low voltage differential EEG activity exhibiting fast oscillatory patterns. NREMS was defined based on the appearance of high-amplitude, low-frequency components in the slow oscillation (SO, 0.75-1.5 Hz), δ (1.5 – 4 Hz) or σ (10 – 15 Hz) frequency range. REMS was recognized based on muscle atonia in combination with the appearance of θ frequency (6 – 8 Hz) in the EEG. Microarousals were scored as maximally 3 consecutive episodes of EEG desynchronization (4 – 12 s) associated with EMG activity that were preceded and followed by NREMS (Franken et al., 1999). The onset of REMS was set as the first epoch with a clearly distinguishable θ peak and the absence of low-frequency activity in the EEG. In case of unipolar recordings, REMS was scored only once low frequencies in the frontal EEG disappeared even when the parietal EEG already showed θ activity. This intermediate sleep-like state was considered as part of NREMS. During CNO injections, NREMS was scored because of a clearly elevated low-frequency component in the EEG and a low-amplitude EMG.

#### Sleep architecture, spectral power composition and dynamics of NREMS*, NREMS and REMS

Based on the visual inspection of the raw traces, we defined a state of NREMS* for experiments involving injection of CNO. NREMS* was scored based on criteria for NREMS, notably the appearance of low-frequency activity and the reduction of muscle tone. NREMS* was also defined to contain the epochs of gradual recovery to NREMS up to the end of the light period in which CNO was injected. No threshold was set to define a point of transition between NREMS* and NREMS. NREMS* was defined for analysis within the first light phase after CNO injection between ZT0 and ZT1.

#### Architectural analyses

We determined NREMS or NREMS* onset latency by calculating the time from the i.p. injection point to the first NREMS or NREMS* epoch (**Figure 5a**). Whenever we compared NaCl or CNO conditions in the first light phase, we refer to NREMS* in the legends. We calculated REMS onset latency as the time difference between the first NREMS or NREMS* epoch and the first sequence of 3 consecutive epochs of REMS (= 12 s) (**Figure 6d**). To compare the amounts of time spent per condition, we calculated the number of epochs per vigilance state as percentage of time within the first 5 h after i.p. injection of NaCl or CNO (**Figure 5b**). The choice of 5 h for this analysis was done because the major VSN stimulation-induced effects occurred during the first 5 h after injection. Furthermore, they permitted inclusion of data from 2 recordings that lasted only 5 h starting from ZT0. The density of microarousals was calculated taking equal times spent in NREMS for NaCl or in NREMS* for CNO conditions (bin = 30 min) (**Figure 5c1**). The same binning was done for the calculation of lengths of NREMS or NREMS* bouts (**Figure 5c2**). For the study of homeostasis, we calculated times spent in either state for 48 h after i.p. injection in hourly bins (**Figure 6**, **Supplementary Figure 6**). The calculation of accumulated times spent in REMS was done in reference to the previous 48 h-baseline recordings (without any injection) in bins of 60 min. Subsequent bins added the accumulated difference from baseline: Accumulated_state_NaCl/CNO (h) = state_NaCl/CNO (h) – state_baseline (h) + Accumulated_state_NaCl/CNO (h-1) (McCarthy et al., 2016). In this plot, data lying on negative slopes indicate a loss of sleep time compared to the baseline, whereas data lying on positive slopes indicate a recovery of sleep times (**Figure 6f**).

#### Spectral analysis

Power spectral analysis was done to identify spectral changes induced by NaCl or CNO exposure according to procedures established in the lab (Fernandez et al., 2018). Relative normalized power spectra were obtained for each condition (NaCl, CNO 1.5 mg/kg, CNO 2.5 mg/kg) by dividing by the summed power values from 0.75 – 35 Hz. We took the first consolidated REMS episode as the timepoint until which we calculated the power spectra starting from the i.p. injection. For comparison of spectra between conditions, a power spectrum after NaCl injection was calculated for every mouse for similar times spent in NREMS, and this separately for both the low and high doses of CNO (**Figure 4a1-a3**). For display, time-frequency plots and dynamics of the characteristic spectral power bands were calculated using Morlet wavelet transforms as described (Osorio-Forero et al., 2021). Total power levels for slow oscillation (SO, 0.75 – 1.5 Hz), δ (1.5 – 4 Hz) or σ (10 – 15 Hz) bands were calculated by summing power values across frequency bins (**Figure 4b**). The dynamics of absolute power in spectral bands of NREMS (δ and σ) were calculated for equal numbers of time bins of NREMS or NREMS* for the NaCl and CNO conditions in non-normalized power spectra. We then normalized power in the δ- and the σ-bands with respect to the corresponding NaCl power bands for the last two time bins of the recording (**Figure 4c1**, **c2**, **8e1**, **e2**). For all these parameters, calculations were done for every recording session first and then means calculated and normalized per condition and finally across animals.

#### Predictive model of homeostatic regulation

The homeostatic process (Process S, see (Franken et al., 2001)) was simulated iteratively based on the 4-s sleep-wake state sequence by assuming that S increases for 4 s that were annotated as wakefulness and REMS and decreases when in NREMS. It is further assumed that the decrease of S during NREMS and NREMS* does not differ. The dynamics of S are tracked by δ power and follow exponential saturating functions with its increase defined as S[t + 1] = UA − (UA − S[t]) × e^−dt/τi^ and its decrease as S[t + 1] = LA + (S[t] − LA) × e^−dt/τd^, where S[t] and S[t + 1] are consecutive values of S delimited by an upper (UA) and lower (LA) asymptote with as time constants τi and τd, for the increase and decrease, respectively. Simulation parameters τi, τd, UA, and LA were taken from (Franken et al., 2006), where they were optimized to fit EEG δ (1 – 4 Hz) dynamics in mice of the same age, sex, and genetic background. The 4 parameters were 7.9 h, 1.9 h, 282 %, and 55 %, respectively. Levels of UA and LA are expressed as % of δ power values obtained at the end of the baseline light periods. The level of S at the onset of the 48-h recording was derived from the simulation assuming steady state at the end of the 48 h (Franken et al., 2001). Process S dynamics were individually determined and averaged over 15 min intervals.

#### Heart rate

Heart rate (HR) was calculated from the RR intervals detected in the EMG in 9/11 animals (Lecci et al., 2017) in bins of 60 min. The first 2 hours were not taken for the HR measures during REMS because there was no REMS (**Figure 2f2**, **Supplementary Figure 6e, f**).

#### Cortical temperature

Temperature was extracted from the voltage measure of the thermistor as indicated above. We calculated all points histograms of temperature values for wake, NREMS and REMS in baseline condition to ensure that the thermistor properly captured the dynamic variation in spontaneous sleep-wake behaviors (Hoekstra et al., 2019). The epoxy thermosensors were partially in contact with the cortical surface, and absolute temperature varied between 28 – 29.5 °C in 24 h-baseline recordings. These values are lower than previously described (Hoekstra et al., 2019). Therefore, only relative temperature changes relative to baseline values just before NaCl or CNO injection were analyzed. We measured the drop of cortical temperature from the i.p. injection moment to the first consolidated REMS period (**Figure 7e**, **f1**). The temperature at which REMS reappeared was determined as the difference between the temperature at the moment of i.p. injection and temperature at the time of the first consolidated REMS episode (**Figure 7f2**).

#### Fiber photometry recordings

The fiber photometry recordings were conducted using a Doric Neuroscience Console (FPC500) that was used to modulate the two LEDs (at 405nm, 475nm) of the of the Doric fluorescence MiniCube LEDs (ilFMC4-G3_IE(400-410)_E(460-490)_F(500-550)_S) at two separate frequencies (208.616 Hz and 333.786 Hz). The combined modulated light was then transmitted through a 400-μm-thick fiberoptic patchcord (MFP_400/430/1100-0.57_1m_FMC-ZF1.25_LAF, Doric Lenses) to the implanted optic canula on the head of the animal. A photodetector integrated into the MiniCube turned the emitted light from the fluorescent sensors into a current signal that was fed back to the Doric Neuroscience Console for acquisition and demodulation. The final sampling rate was 60 Hz. The measured power at the tip of the optic fiber was 12.5 µW for isosbestic 405 nm excitation and 25 µW for the 475 nm. For the green emission (jGCaMP8s), the ΔF/F0 signal was computed using, as F0, the fitted isosbestic Ca^2+^-independent signal (from the violet excitation LED) (**Figure 3c**) (Osorio-Forero et al., 2024).

#### Quantification of z-scored LC ΔF/F0 for NaCl and CNO (1.5 mg/kg) injections relative to pre-injection activity

The mean z-scored ΔF/F0 fiber photometry signal, calculated over 1 h after the first scored NREMS or NREMS* epoch following injection of NaCl or CNO, was divided by the mean z-scored ΔF/F0 fiber photometry signal for 1 h before the injection (**Figure 3d1**). For the quantification of the LC activity after partial recovery from CNO effects, we calculated the mean z-scored ΔF/F0 fiber photometry signal for 1 h after the first scored REM epoch (**Figure 3d2**).

#### Quantification of the spectral composition of the LC ΔF/F0 for NaCl and CNO (1.5 mg/kg) injections compared to pre-injection

We computed the Gabor-Morlet Wavelet Transform of the ΔF/F0 fiber photometry signal using a custom-made code in Python in a frequency range between 0.001 and 0.251 Hz with a step of 0.005 Hz. We then calculated the ratio between the power at the “lower” spectral components (0.01 – 0.03 Hz) and the “higher” spectral components (0.1 – 0.25 Hz) for time points similar to the ones for mean z-scored ΔF/F0 (**Figure 3d3**, **d4**, **Supplementary Figure 3**).

### Statistical analysis

Statistical tests were done using MATLAB (Mathworks) or R (V.4.1.2., The R Foundation for Statistical Computing). In all figures, the final and significant p values resulting from these tests are written directly in the figure panels. The n for the number of animals used is also indicated in each panel. In Figure 1, the n number corresponds to the number of cells that were recorded. For all figure panels, the statistical tests used are indicated in the legends. To select the appropriate test, datasets were tested for normality via the Shapiro-Wilk test and for homogeneity of variances using the Bartlett test. Then, we performed parametric one-way (repeated-measures) ANOVA or (non-parametric) Friedmann tests, followed by *post hoc* tests (paired t-tests or Wilcoxon signed-rank or rank sum tests). Bonferroni correction was used to correct for multiple comparisons. For the correlation analysis, we used a linear regression model (fitlm in Matlab). All box-and-whisker plots represent data as done by default in Matlab: the central mark indicates the median, and the bottom and top edges of the box indicate the 25^th^ and 75^th^ percentiles, respectively. The whiskers extend to the most extreme data points not considered outliers, and the outliers are plotted individually using the ’+’ marker symbol. In the presence of the outliers, the whiskers are given as ± 1.5 times the interquartile range. Complete information for all statistical tests is indicated in the **Supplementary Table 1** for all Figure panels.

## Supplementary Figures and Legends

**Supplementary Figure 1.**
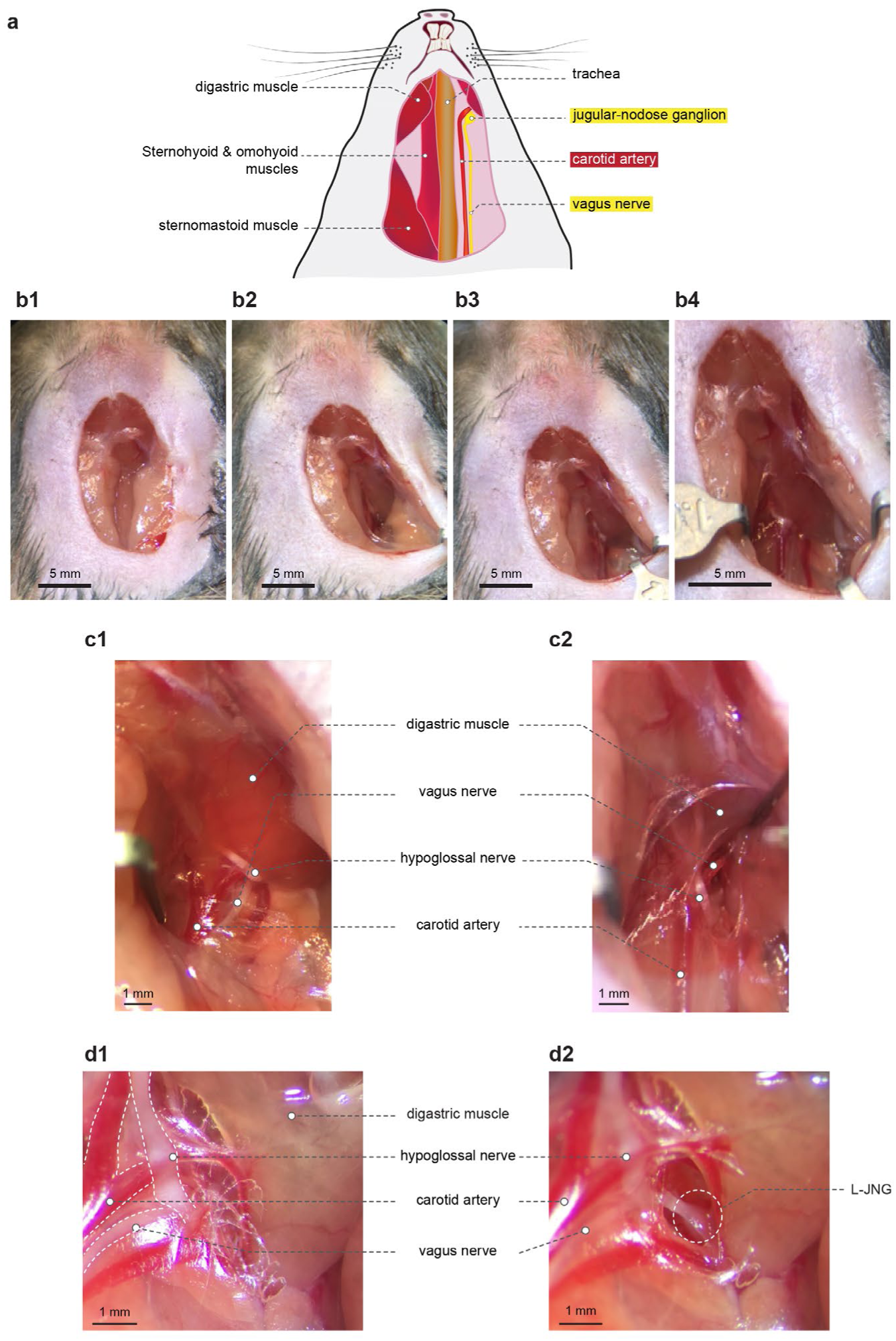
Illustration of surgery to access the L-JNG for viral injection. a. Schematic of major anatomical elements visible upon accessing the throat area over the L-JNG. b. Early steps of the surgery. Access to the ganglion starts by gently displacing the salivary glands (b1, b2) and via putting retractors on the ipsilateral sternomastoid (b3) and the contralateral omohyoid (b4) muscles. c. Two views (c1, c2) of the exposure of both vagal and hypoglossal nerves in the late steps of the surgery. d. Two zoom-in views of the site of the L-JNG location before (d1) and after (d2) removing connective tissue.

**Supplementary Figure 2.**
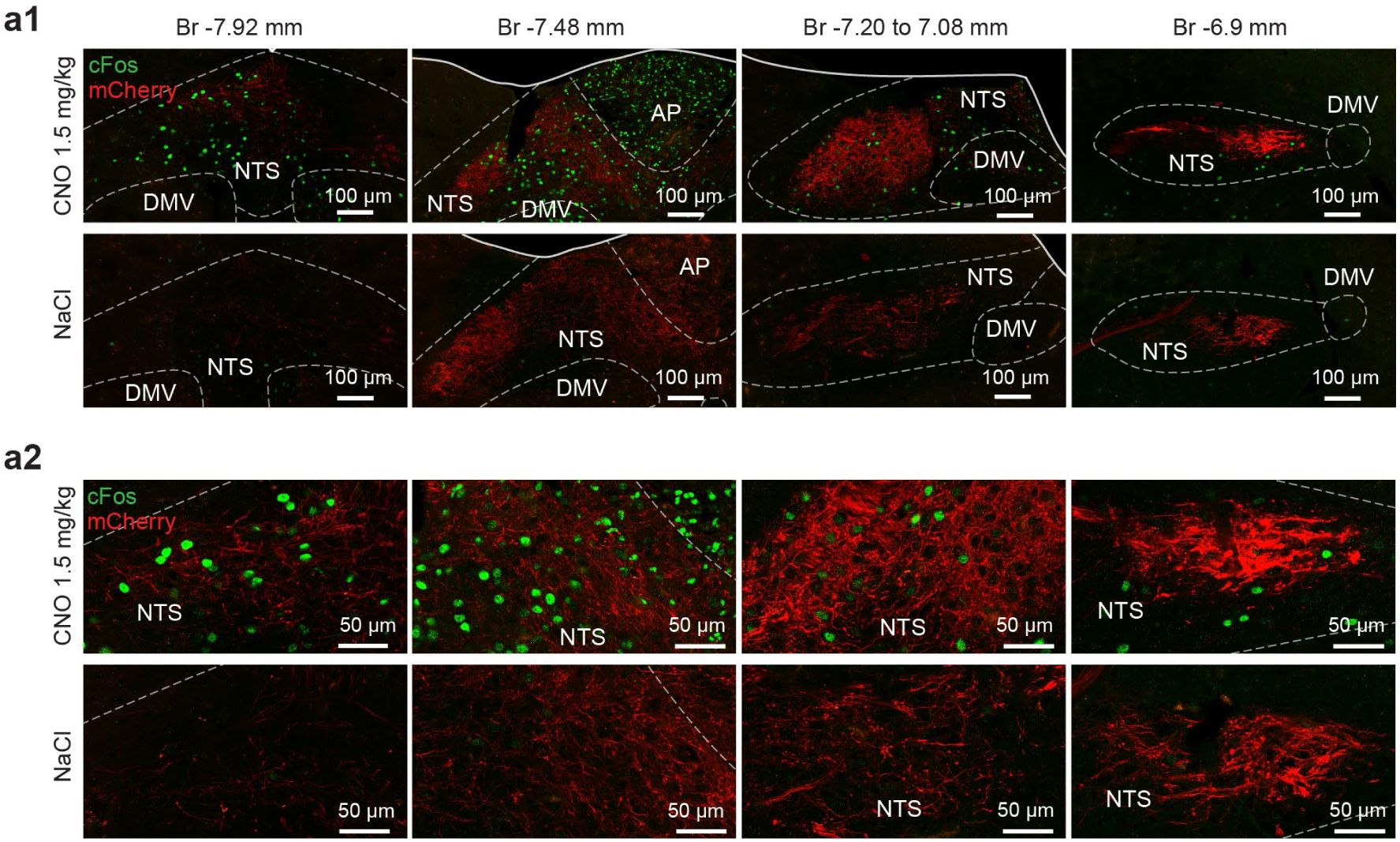
Further illustration of cFos expression in vagal-recipient areas of the dorsal vagal complex. a1, a2. Zoom-out (a1) and Zoom-in (a2) of fluorescent micrographs shown in **Figure 2c1**, plus an additional one for Br -6.9 mm, for CNO (1.5 mg/kg, top row) and NaCl (bottom row) injections.

**Supplementary Figure 3.**
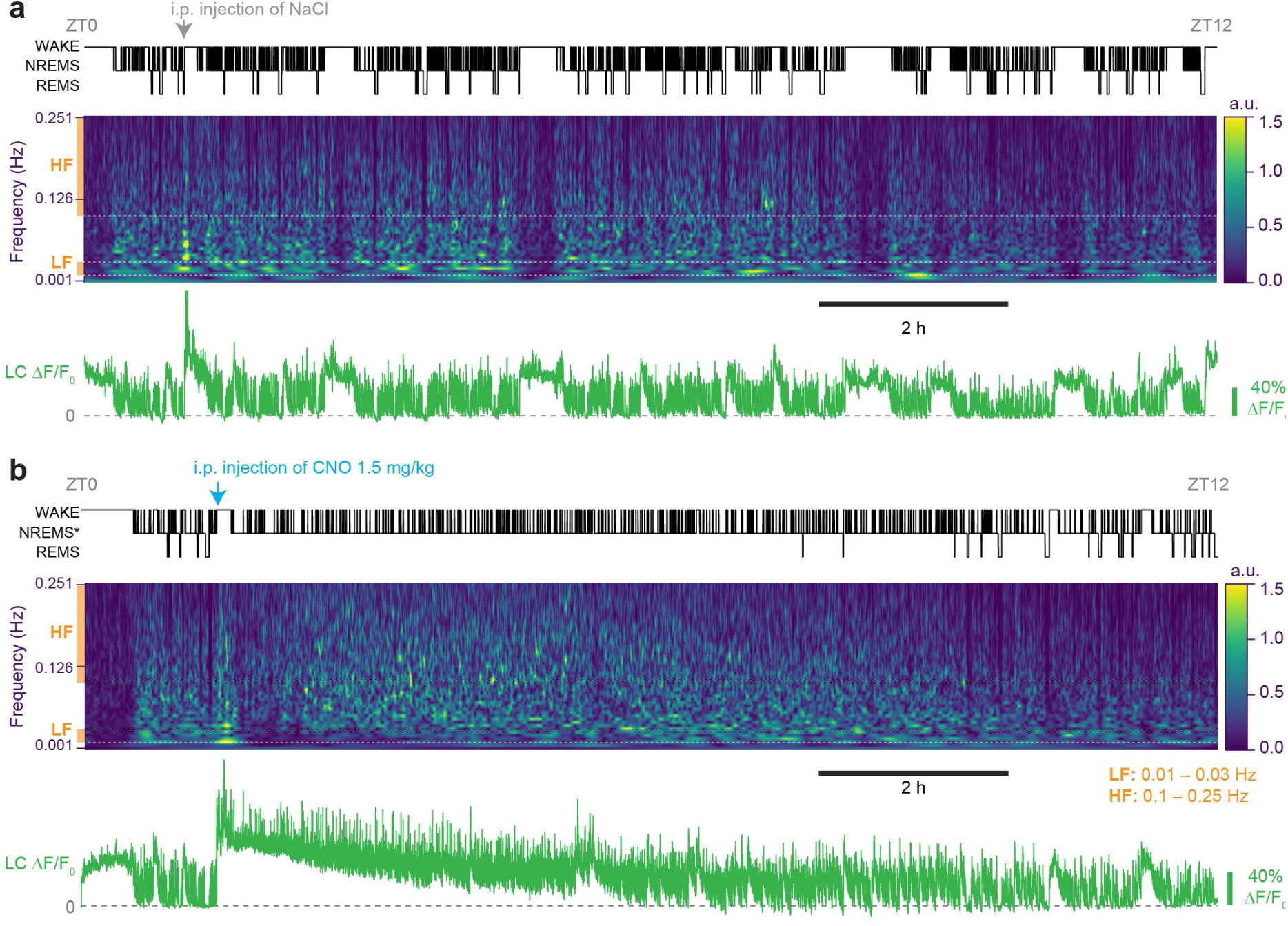
Spectrograms of LC activity, with indication of high- and low-frequency components. a. Data from **Figure 3c1**, replotted together with spectrogram. Note the disappearance of low-frequency activity (LF, 0.01-0.03 Hz, orange bar) after chemogenetic VSN stimulation. HF, high-frequency activity (0.1-0.25 Hz), which was used to calculate the spectral power ratios calculated in **Figure 3d3**,**d4**. b. As a, for data from **Figure 3c2**.

**Supplementary Figure 4.**
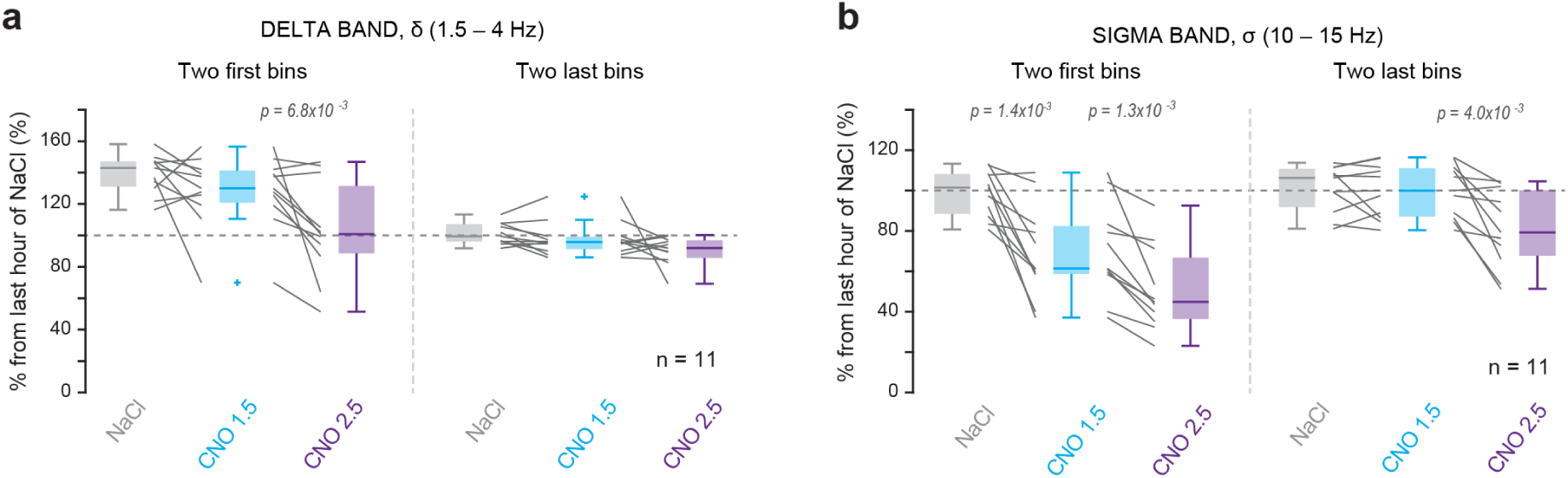
Quantification of absolute power dynamics in response to vagal sensory stimulation. a. Quantification and statistical analysis for data presented in **Figure 4c1**, with mean absolute power levels (in % of NaCl values) shown for the first and the last two time bins for δ power. Friedman or one-way RM-ANOVA tests, with post hoc Wilxocon signed-rank or paired t-tests, with Bonferroni-corrected p=0.0125. Significant p values are indicated. b. As a, for σ power shown in **Figure 4c2**. One-way repeated-measures ANOVA with post hoc t-tests, with Bonferroni-corrected p=0.0125. Significant p values are shown.

**Supplementary Figure 5.**
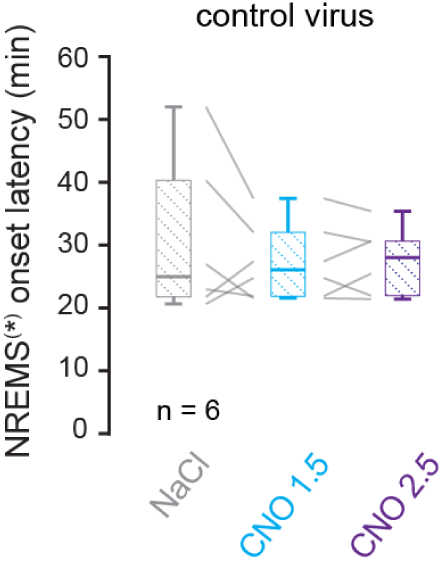
Quantification of NREMS and NREMS* onset latencies in control animals. Box-and-whisker plots of NREMS and NREMS* (referred to together as NREMS^(^*^)^ in y-axis) onset latency for animals expressing a non-excitatory Dreadd-expressing viral construct (see Methods, *Surgeries for viral injections in L-JNG, Expression of Dreadd-unrelated constructs to control for CNO effects*). Control data for Figure 5a. One-way repeated-measures ANOVA with factor ‘Treatment’, with post hoc t-tests. No significance was obtained.

**Supplementary Figure 6.**
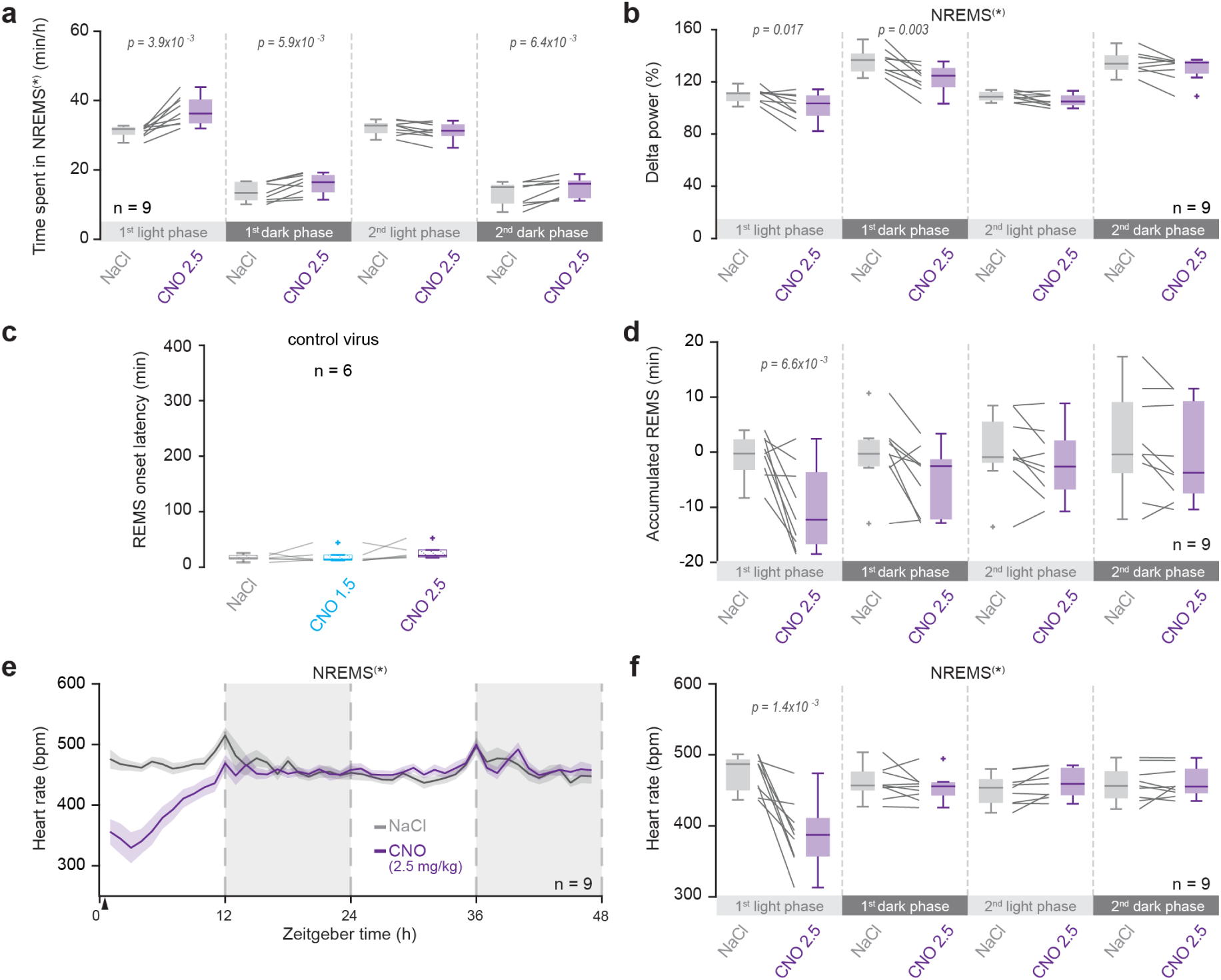
Quantification of heart rate and REMS timing for two light and dark phases after VSN stimulation (between ZT0 and ZT1 in the first light phases). a. Box-and-whisker plots for times spent in NREMS* or NREMS (referred to together as NREMS^(^*^)^ in y-axis) in min/h, as shown in Figure 6a. Student paired t-tests. b. As a, for mean δ power values shown in Figure 6b. Wilcoxon signed-rank tests or Student paired t-tests with Bonferroni correction for multiple comparison (p = 0.0125). Significant p values are indicated. c. Box-and-whisker plots for REMS onset latencies for control virus-expressing animals, complementing the data shown in Figure 6d. Friedman test with factor ‘Treatment’, not significant. d. Box-and-whisker plots for accumulated REMS time, from data shown in Figure 6e. Paired t-tests with Bonferroni-corrected p = 0.0125. Significant p values are indicated. e. Time course of heart rate for NREMS* or NREMS over 2 light and dark phases, following CNO or NaCL injection between ZT0-ZT1. Arrowhead denotes the time point of injection. f. Box-and-whisker plots for mean heart rates across the light and dark phases shown in panel e Paired t-tests, with Bonferroni-corrected p = 0.0125, significant p values are given.

**Supplementary Figure 7.**
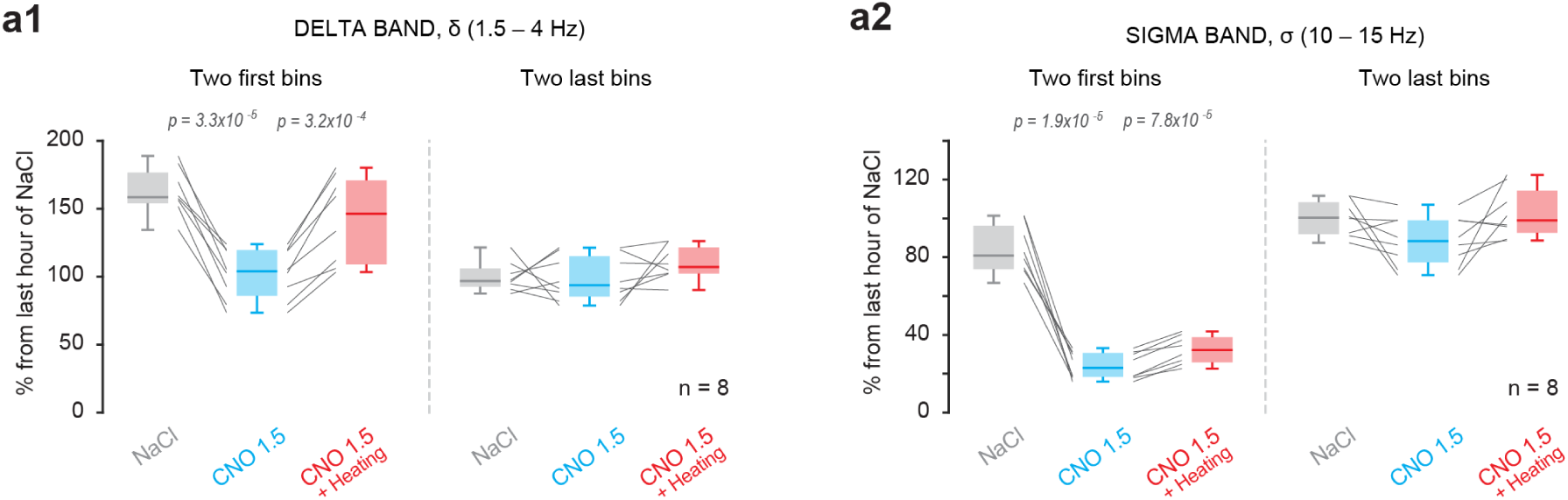
Quantification of data obtained from ambient heating during vagal sensory stimulation. Quantification of data presented in Figure 8e, with mean power levels shown for the first two and the last two-time bins for δ (a1) and σ (a2) power. One-way repeated-measures ANOVA with factor ‘Treatment’, with post hoc t-tests. Bonferroni-corrected p = 0.0125. Significant p values are indicated.

**Supplementary Table 1.** Summary of statistical analyses presented in Main and Extended Data Figures. This table describes statistical tests used for each figure/panel in this paper. Per figure panel, the number of animals, the test statistics (F/t values) and the degrees of freedom (Df/df) are indicated. Post hoc p values are given in a separate column and only for significant results. Bonferroni-corrected p values are given in the legends.

| Figs and panels | Type of experiment | Tests used | n numbers (animals unless otherwise indicated) | Test statistics (F, t) & degrees of freedom (Df, df) | P values | Post hoc tests with multiple comparisons, Test statistics & df | Post-hoc tests, p values |
| --- | --- | --- | --- | --- | --- | --- | --- |
| **Fig 2c2** | cFos-expression in dorsal vagal complex after NaCl or CNO inj. | Unpaired t-test | For NaCl: n = 7, for CNO: n= 8 | For NTS left: NaCl vs CNO1.5: t= 3.09, df=7<br>For NTS right: NaCl vs CNO1.5: t= 4.32, df=7<br>For AP: NaCl vs CNO1.2: t=3.61, df=7 | For NTS left: p= 0.017<br>For NTS right: p= 3.1*10<sup>-3</sup><br>For AP: p= 8.6*10<sup>-3</sup> | *none* |  |
| **Fig 2f3** | Heart rate changes for NREMS and REMS after NaCl or CNO (1.5 mg/kg) inj. | Wilcoxon signed-rank test for NREMS<br>Paired t-test for REMS | n = 9 | For NREMS: NaCl vs CNO1.5: V=45<br>For REMS: NaCl vs CNO1.5: t=6.4753, df=8 | For NREMS: NaCl vs CNO1.5: p=0.0039<br>For REMS: NaCl vs CNO1.5: p= 1.9*10<sup>-4</sup> | *none* |  |
| **Fig 2h** | Mean distance moved after NaCl or CNO (1.5 mg/kg) inj. | Paired t-test | n = 5 | For NaCl vs CNO1.5: t=-3.18, df=4 | For NaCl vs CNO1.5: p=0.033 | *none* |  |
| **Figure 3a2** | cFos expression levels | For LC left, Wilcoxon-Mann- | n = 6 for NaCl, n = | For LC left: U=34 | For LC left: p = 0.0087 | *none* |  |
|  | in LC after NaCl or CNO (1.5 mg/kg) inj. | Whitney two-sample rank test<br>For LC right: Wilcoxon-Mann-Whitney two-sample rank test | 6 for CNO | For LC right, U=33 | For LC right: P = 0.015 |  |  |
| **Figure 3d1** | LC population activity magnitude after NaCl or CNO (1.5 mg/kg) inj | Paired t-test | n = 5 | For NaCl vs CNO:<br>t=-5.24, df=4 | For NaCl vs CNO:<br>p = 0.0063 | none |  |
| **Figure 3d2** | LC population activity magnitude after CNO (1.5 mg/kg) inj or after recovery | Paired t-test | n = 5 | For NaCl vs CNO:<br>t=-10.39 df=4 | For NaCl vs CNO:<br>p = 0.00048 | none |  |
| **Figure 3d3** | LC population activity spectrogram composition after NaCl or CNO (1.5 mg/kg) inj | Paired t-test | n = 5 | For NaCl vs CNO:<br>t=9.09, df=4 | For NaCl vs CNO:<br>p = 0.00081 | none |  |
| **Figure 3d4** | LC population activity spectrogram composition after CNO (1.5 mg/kg) inj or after recovery | Paired t-test | n = 5 | For NaCl vs CNO:<br>t=-10.20, df=4 | For NaCl vs CNO:<br>p = 0.00052 | none |  |
| **Fig 4b** | Power in bands after NaCl or CNO (1.5 or 2.5 mg/kg) inj. until first REMS episode | Paired t-test | n = 11 | For NaCl vs CNO 1.5:<br>SO: t=-3.26, df=10<br>// $\delta$: t=-4.41, df=10<br>// $\sigma$: t=4.71, df=10<br>For NaCl vs CNO 2.5:<br>SO: t=-3.85, df=10<br>// $\delta$: t=-4.04, df=10<br>// $\sigma$: t=6.61, df=10 | For NaCl vs CNO 1.5:<br>SO: p=8.5*10<sup>-3</sup><br>// $\delta$: p=1.3*10<sup>-3</sup><br>// $\sigma$: p=8.3*10<sup>-4</sup><br>For NaCl vs CNO 2.5:<br>SO: p=3.2*10<sup>-3</sup><br>// $\delta$: p=2.3*10<sup>-3</sup><br>// $\sigma$: p=6.0*10<sup>-5</sup> | none | |
| **Fig 5a** | NREMS* and NREMS onset latency after NaCl or CNO (1.5 mg/kg or 2.5 mg/kg) inj. | One-way ANOVA with factor 'Treatment', followed by Student's paired t-test | n = 11 | For treatment: F<sub>2,30</sub>=28.15 | For treatment: p=1.3*10<sup>-7</sup> | For NaCl vs CNO1.5: t = 5.35, df = 10<br>For NaCl vs CNO2.5: t = 1.69, df = 10 | For NaCl vs CNO1.5: p = 3.2*10<sup>-4</sup><br>For CNO1.5 vs CNO2.5: p = 0.12 |
| **Fig 5b** | Times spent in Wake, NREMS, NREMS* and REMS after NaCl or CNO (1.5 mg/kg or 2.5 mg/kg) inj. | For Wake: One-way ANOVA with factor 'Treatment', followed by Student's paired t-test<br>For NREMS(*), REMS: | n = 11 | For WAKE: F<sub>2,30</sub>=36.6 | For WAKE: p = 8.9*10<sup>-9</sup><br>For NREMS(*): | WAKE: For NaCl vs CNO 1.5: t=8.84, df=10<br>For CNO 1.5 vs CNO 2.5: t=3.62, df=10<br>NREMS(*): | WAKE: For NaCl vs CNO 1.5: p = 9.8*10<sup>-4</sup><br>For CNO 1.5 vs CNO 2.5: p = 9.8*10<sup>-4</sup><br>NREMS(*): |
| | | Friedman test with factor 'Treatment', followed by Wilcoxon signed-rank test | | For NREMS(*):<br>Chisq=22, DF1=2, DF2=20<br>For NREMS(*):<br>Chisq=14.42, DF1=2, DF2=20 | $p = 1.7 \times 10^{-5}$<br>For REMS:<br>$p = 7.6 \times 10^{-4}$ | For NaCl vs CNO 1.5:<br>V=0<br>For CNO 1.5 vs CNO 2.5:<br>V=0<br>REMS:<br>For NaCl vs CNO 1.5:<br>V=46<br>For CNO 1.5 vs CNO 2.5:<br>V=66 | For NaCl vs CNO 1.5: $p = 9.8 \times 10^{-4}$<br>For CNO 1.5 vs CNO 2.5: $p = 9.8 \times 10^{-4}$<br>REMS:<br>For NaCl vs CNO 1.5: $p = 0.28$<br>For CNO 1.5 vs CNO 2.5: $9.8 \times 10^{-4}$ |
| **Fig 5c1** | Microarousal density in NREMS and NREMS* after NaCl or CNO (1.5 mg/kg or 2.5 mg/kg) inj. | Friedman test with factor 'Treatment', followed by Wilcoxon signed rank test | n = 11 | For treatment for 1<sup>st</sup> 30-min period:<br>Chisqr=6.8<br>Df1=2, Df2=20<br>For 2<sup>nd</sup> 30-min period:<br>F=0.42, Df1=2, Df2=30 | For treatment for 1<sup>st</sup> 30-min period:<br>$p = 0.032$<br>For 2<sup>nd</sup> 30-min period:<br>$p = 0.078$ | For NaCl vs CNO 1.5:<br>V=45<br>For CNO 1.5 vs CNO 2.5:<br>V=5.5 | For NaCl vs CNO 1.5: $p = 0.32$<br>For CNO 1.5 vs CNO 2.5: $p = 0.021$ |
| **Figure 5c2** | Bout lengths of NREMS and NREMS* after NaCl or CNO (1.5 mg/kg or 2.5 mg/kg) inj. | For treatment for 1<sup>st</sup> 30-min period: One-way ANOVA with factor 'Treatment'<br>For treatment for 2<sup>nd</sup> 30-min period: Friedman test with factor 'Treatment', | n = 11 | For treatment for 1<sup>st</sup> 30-min period:<br>F=0.63, df1=2, df2=30<br>For treatment for 2<sup>nd</sup> 30-min period:<br>Chisqr=4.90, Df1=2, Df2=20 | For treatment for 1<sup>st</sup> 30-min period:<br>$p = 0.54$<br>For treatment for 2<sup>nd</sup> 30-min period:<br>$p = 0.086$ | none | |
| **Fig 6d** | Latency to REMS onset after NaCl or CNO (1.5 or 2.5 mg/kg) inj | Friedman test with factor 'Treatment', followed by post hoc Wilcoxon signed-rank tests | n = 11 | For treatment: Chisqr=20.18<br>Df1=2, DF2=20 | For treatment: $p = 4.1 \times 10^{-5}$ | For NaCl vs CNO 1.5: V=2<br>For CNO 1.5 vs CNO 2.5: V=0 | For NaCl vs CNO 1.5: $p = 2.9 \times 10^{-3}$<br>For CNO 1.5 vs CNO 2.5: $p = 9.8 \times 10^{-4}$ |
| **Fig 7c** | Cortical temperature measures in baseline for wake, NREMS, REMS | One-way ANOVA with factor 'Vigilance state', followed by Student's paired t-test | n = 7 | For Vigilance state: F=34.98, Df1=1, Df2=6 | For Vigilance state: $p = 0.001$ | For Wake vs NREMS: $t=6.92$, $df=6$<br>For NREMS vs REMS: $t=-4.74$, $df=6$ | For Wake vs NREMS: $p = 4.5 \times 10^{-4}$<br>For NREMS vs REMS: $p = 3.2 \times 10^{-3}$ |
| **Figure 7e** | Cortical temperature drop after NaCl or CNO (1.5 or 2.5 mg/kg) inj | Friedmann test with factor 'Treatment', followed by post hoc Wilcoxon signed-rank test | n = 7 | For Treatment: Chisq=11.14<br>Df1=2, Df2=12 | For Treatment: $p = 0.0038$ | For NaCl vs CNO 1.5: V = 28 | For NaCl vs CNO 1.5: $p = 0.016$ |
| **Figure 7f1, f2** | Linear correlation between REMS onset latency and temperature drops | Linear Regression | For panel f1: n = 14<br>For panel f2: n = 14 | For regression in panel f1: F= 204, $df = 12$<br>For regression in panel f2: F= 16.2, $df = 12$ | For regression in panel f1: $p = 6.7 \times 10^{-9}$<br>For regression in panel f2: $p = 0.0017$ | | |
| **Figure 7g** | Rectal temperature drop after NaCl or CNO (1.5 mg/kg) | Paired t-test | n = 7 | For NaCl vs CNO 1.5, time point 1: $t=4.55$, $df=6$ | For NaCl vs CNO 1.5, time point 1: $p = 3.9 \times 10^{-3}$ | none | none |
| | | | | For NaCl vs CNO 1.5, time point 2: t= 4.41, df= 6 | For NaCl vs CNO 1.5, time point 2: p = $4.5 \times 10^{-3}$ | | |
| **Figure 8c** | Cortical temperature drop after NaCl or CNO (1.5 mg/kg) inj +/- heating | Friedmann test for factor 'treatment', followed by post hoc Wilcoxon | n = 8 | For treatment: Chisq=16.0 Df1=2, Df2=14 | For treatment p=0.00033 | For NaCl vs CNO-heating: V=36<br>For CNO - vs CNO + heating: V=0 | For NaCl vs CNO-heating: p = $7.8 \times 10^{-3}$<br>For CNO- vs CNO + heating: p = $7.8 \times 10^{-3}$ |
| **Figure 8d** | Latency to REMS onset after CNO (1.5 mg/kg) inj +/- heating | Friedmann test for factor 'treatment', followed by post hoc Wilcoxon | n = 8 | For treatment: Chisq=14.25 Df1=2, Df2=14 | For treatment p=0.0008 | For NaCl vs CNO-heating: V=0<br>For CNO- vs CNO + heating: V=35 | For NaCl vs CNO-heating: p = $7.8 \times 10^{-3}$<br>For CNO- vs CNO + heating: p = 0.016 |
| **Figure 8f1** | Linear correlation between REMS onset latency and temperature drops | Linear Regression | n = 16 | For regression: F= 103, df = 14 | p = $7.7 \times 10^{-8}$ | none | |
| **Figure 8f2** | Linear correlation between REMS onset latency and temperature at REMS onset | Linear Regression | n = 16 | For regression: F= 3.65, df = 14 | p = 0.08 | none |  |
| **Fig S4a** | Power in $\delta$ band | Friedmann or RM- | n =11 | For treatment, | For treatment, first two bins: | For first two bins: | For first two bins: |
|  | relative to last hour (hour 5) of NaCl for NaCl or CNO (1.5 or 2.5 mg/kg) inj | ANOVA with factor 'Treatment', followed by post hoc Wilcoxon or Student t-tests |  | first two bins:<br>Chisqr=12.18, df1=2, df2=20<br>For treatment, last two bins:<br>F=3.97, df1=2, df2=30 | p=0.0023<br>For treatment, last two bins: p=0.030 | For NaCl vs CNO1.5: V=49<br>For CNO1.5 vs CNO2.5: V=62<br>For last two bins:<br>For NaCl vs CNO1.5: t=1.36, df=10<br>For CNO1.5 vs CNO2.5: t=1.42, df=10 | For NaCl vs CNO1.5: p=0.17<br>For CNO1.5 vs CNO2.5: p=0.0068<br>For last two bins:<br>For NaCl vs CNO1.5: p=0.20<br>For CNO1.5 vs CNO2.5: p=0.19 |
| **Fig S4b** | As Fig S4a, for power in $\sigma$ band | RM-ANOVA Factor treatment followed by post hoc Student t-test | n =11 | For treatment, first two bins:<br>F=17.58, df1=2, df2=30<br>For treatment, last two bins:<br>F=5.97, df1=2, df2=30 | For treatment, first two bins: p=8.8*10<sup>-6</sup><br>For treatment, last two bins: p= 0.0066 | For first two bins:<br>For NaCl vs CNO1.5: t=4.36, df=10<br>For CNO1.5 vs CNO2.5: t=4.41, df=10<br>For last two bins:<br>For NaCl vs CNO1.5: t=0.41, df=10<br>For CNO1.5 vs CNO2.5: t=3.72, df=10 | For first two bins:<br>For NaCl vs CNO1.5: p=0.0014<br>For CNO1.5 vs CNO2.5: p=0.0013<br>For last two bins:<br>For NaCl vs CNO1.5: p=0.69<br>For CNO1.5 vs CNO2.5: p=0.0040 |
| **Fig S5** | Delay to NREMS or NREMS* onset, control | RM-ANOVA Factor treatment followed by post hoc | n = 6 | For treatment: F=0.27, Df1=2, Df2=15 | For treatment: P=0.76 | none |  |
|  | experiments | Student t-test |  |  |  |  |  |
| **FigS6a** | Time spent in NREMS or NREMS* | Wilcoxon signed-rank or t-tests | n = 9 | For NaCl vs CNO:<br>1<sup>st</sup> light phase: V=0<br>1<sup>st</sup> dark phase: t= -3.72, df = 8<br>2<sup>nd</sup> light phase: t= -3.66, df = 8<br>2<sup>nd</sup> dark phase: t= 1.85, df = 8 | For NaCl vs CNO:<br>1<sup>st</sup> light phase: p = 0.0039<br>1<sup>st</sup> dark phase: p = 5.9 *10<sup>-3</sup><br>2<sup>nd</sup> light phase: p = 0.11<br>2<sup>nd</sup> dark phase: p = 6.4 *10<sup>-3</sup> | none |  |
| **Fig S6b** | Delta power changes after CNO (2.5 mg/kg) injection in % of NaCl across two light and dark phases | Paired t-test | n = 9 | For NaCl vs CNO:<br>1<sup>st</sup> light phase: t=2.98, df=8<br>1<sup>st</sup> dark phase: t= 4.21, df = 8 | For NaCl vs CNO:<br>1<sup>st</sup> light phase: p = 0.017<br>1<sup>st</sup> dark phase: p = 2.9 *10<sup>-3</sup> | none |  |
| **Fig S6c** | Delay to REMS onset after NaCl or CNO (1.5 mg/kg or 2.5 mg/kg) inj. in mice expressing a non- | Friedmann test followed by post hoc Wilcoxon signed rank test | n = 6 | For treatment: Chisqr=4.33, df1=2, df2=15 | For treatment: p=0.11 | none |  |
|  | Dread-related control virus |  |  |  |  |  |  |
| **Fig S6d** | Accumulated REMS time over 48 h for NaCl or CNO (2.5 mg/kg) injection | Paired t-tests | n = 9 | For NaCl vs CNO:<br>1<sup>st</sup> light phase: t= 3.66, df = 8<br>1<sup>st</sup> dark phase: t= 2.78, df = 8<br>2<sup>nd</sup> light phase: t= 1.85, df = 8<br>2<sup>nd</sup> dark phase: t= 1.85, df = 8 | For NaCl vs CNO:<br>1<sup>st</sup> light phase: p=0.0064<br>1<sup>st</sup> dark phase: p=0.024<br>2<sup>nd</sup> light phase: p=0.1<br>2<sup>nd</sup> dark phase: p=0.1 | *none* |  |
| **Fig S6f** | Heart rate changes over 48 h after NaCl or CNO (2.5 mg/kg) inj. For NREMS*, NREMS | Paired t-tests | n = 9 | For NREMS*:<br>For NaCl vs CNO, 1<sup>st</sup> light phase: t=4.75, df=8<br>1<sup>st</sup> dark phase: t=1.1, df=8<br>2<sup>nd</sup> light phase: t=-2.40, df=8<br>2<sup>nd</sup> dark phase: t=-0.49, df=8 | For NREMS*:<br>For NaCl vs CNO, 1<sup>st</sup> light phase: p = 1.4*10<sup>-3</sup><br>1<sup>st</sup> dark phase: p = 0.3<br>2<sup>nd</sup> light phase: p = 0.043<br>2<sup>nd</sup> dark phase: p = 0.63 | *none* |  |
| **Figure S8a1** | Mean Power in $\delta$ band after NaCl or CNO (1.5 mg/kg) inj +/- heating | RM-ANOVA, Factor 'Treatment', followed by post hoc paired t-tests | n =8 | For treatment, first two bins: F= 13.4, Df1=2, Df2=8<br>For treatment, last two bins: F= 1.6, Df1=2, Df2=8 | For treatment, first two bins: p = $1.8 \times 10^{-4}$<br>For treatment, last two bins: p = 0.22 | For NaCl vs CNO: t=9.36, df=7<br>For CNO - vs CNO + heating: t=-6.55, df=7<br>none | For NaCl vs CNO: p=$3.3 \times 10^{-5}$<br>For CNO - vs CNO + heating: p=$3.2 \times 10^{-4}$<br>none |
| **Figure S8a2** | As Fig S8a1, for Power in $\sigma$ band | RM-ANOVA, With factor 'Treatment', followed by post hoc paired t-tests | n =8 | For treatment, first two bins: F= 93.2 Df1=2, Df2=8<br>For treatment, last two bins: F= 3.34, Df1=2, Df2=8 | For treatment, first two bins: p = $3.6 \times 10^{-11}$<br>For treatment, last two bins: p = 0.055 | For NaCl vs CNO: t=10.1, df=7<br>For CNO - vs CNO + heating: t=-8.18, df=7<br>none | For NaCl vs CNO: p=$1.9 \times 10^{-5}$<br>For CNO - vs CNO + heating: p=$7.9 \times 10^{-5}$<br>none |

